# Chloroplast genome editing of Rubisco boosts photosynthesis and plant growth

**DOI:** 10.1101/2025.01.02.631008

**Authors:** Wataru Yamori, Issei Nakazato, Qu Yuchen, Yukina Sanga, Tomoko Miyata, Ryo Uehara, Yuma Noto, Keiichi Namba, Hiroshi Fukayama, Hiroyoshi Matsumura, Shin-ichi Arimura

**Author notes:** These authors contributed equally (Wataru Yamori, Issei Nakazato).

## Abstract

Photosynthetic inefficiencies limit the productivity and sustainability of crop production and the resilience of agriculture to future societal and environmental challenges. Ribulose-1,5-bisphosphate carboxylase/oxygenase (Rubisco) has inherently low catalytic efficiency, making it a key target for photosynthesis and crop improvement. However, introducing mutations to the chloroplast-encoded Rubisco large subunit (*rbc*L), which contains the enzyme’s catalytic sites, is technically challenging. In this study, we successfully generated a range of chloroplast-genome-edited *Arabidopsis thaliana* plants targeting *rbcL* by a targeted base editor, ptpTALECD. The M309I and D397N substitutions in *rbcL* resulted in an increased Rubisco catalytic rate (*k*_cat_) without any reductions of Rubisco content, thereby enhancing photosynthetic rates and plant growth under both current atmospheric CO_2_ concentrations (i.e., 381 μmol mol^−1^) and projected future concentrations (i.e., 549 μmol mol^−1^). Cryo-electron microscopy (cryo-EM) structural analysis showed that the M309I and D397N substitutions, although located far from the catalytic site, induce structural alterations in the catalytic (60s) loops. Our findings highlight the potential of Rubisco engineering to improve plant photosynthesis and growth, and underscore the unique opportunities that chloroplast genome editing offers for enhancing photosynthesis and crop productivity and reducing atmospheric CO_2_ levels in a non-GMO context.

## Main text

Climate change has been accelerating, with rising atmospheric CO_2_ levels contributing to higher global temperatures since the Industrial Revolution^1,2^ and exacerbating the food crisis. Global food production is projected to need to increase by more than 30% by 2050 to meet growing demands^3,4^. Plants, as the foundation of the global food supply, are susceptible to climate change and play a crucial role in regulating atmospheric CO_2_. Rubisco is inefficient, making its catalytic reactions the limiting step in photosynthetic CO_2_ fixation under physiologically relevant conditions. Engineering Rubisco with a high carboxylation rate into plant chloroplasts is a promising approach to increasing global crop yields and reducing atmospheric CO_2_ levels^4–6^. Mathematical modeling suggests that introducing Rubisco with a high carboxylation rate could potentially result in over a 25% increase in crop yields^7,8^.

Engineering Rubisco in vascular plants has traditionally been conducted exclusively in tobacco (*Nicotiana tabacum*), where well-established procedures for chloroplast transformation, the foreign gene insertion, are available^9,10^. Rubisco is a hexadecameric complex composed of eight large subunits encoded by the *rbcL* chloroplast genome-encoded gene and eight small subunits encoded by a family of *rbcS* nuclear genes. Efforts have been made to identify new Rubisco variants with higher turnover rates from diverse natural species or to develop hybrid Rubisco enzymes to replace the endogenous plant Rubisco^6,8,11–18^. However, attempts to engineer vascular plants with these Rubisco variants have largely been unsuccessful, at best growing slower than wild type even in enhanced 1% (= 10,000 μmol mol^−1^) CO_2_ conditions, primarily due to issues with inefficient assembly and poor solubility of heterologously expressed Rubisco. These methods, even if succeeded, would pose another hurdle of GM regulation in many countries, so here, we tried to carry out genome editing also for future social acceptance.

In recent years, directed evolution approaches using Rubisco-dependent *Escherichia coli* selection^19^ and evolutionary studies of Rubisco from C3 to C4 plants of the genus *Flaveria*^20^ have identified several candidate residues in RbcL that could potentially improve Rubiscos carboxylation rate. However, how the structural changes within Rubisco enhance its characteristics remains still unexplored. The next challenge is to introduce these residues directly into different plant species under the natural expression levels. In this study, we successfully generated genome-edited *A. thaliana* lines with stable point mutations in *rbcL* to assess their effects on Rubisco function/structure, photosynthesis, and plant growth.

### Amino acid modification in *RbcL*

We initially focused on six amino acid substitutions in the Rubisco large subunit (RbcL) that have been suggested to enhance Rubisco’s carboxylation efficiency [M309I^20^, L74M, I393M, D397N, A414T, and P415A^19^], as shown in Fig. 1. Of these, M309I, D397N, and A414T could be introduced by C:G-to-T:A base editing^21^. By this base editing, L74 and P415 could be substituted for similar amino acids, Phe and Ser, respectively. The residues L74, D397, A414, and P415 are conserved across all 32 investigated crop and tree species, while M309 is conserved in 26 of these species (Table S1). Consequently, we attempted to introduce a total of five mutations into *A. thaliana*: L74F, M309I, D397N, A414T, and P415S (Fig. 1b).

**Fig. 1.**
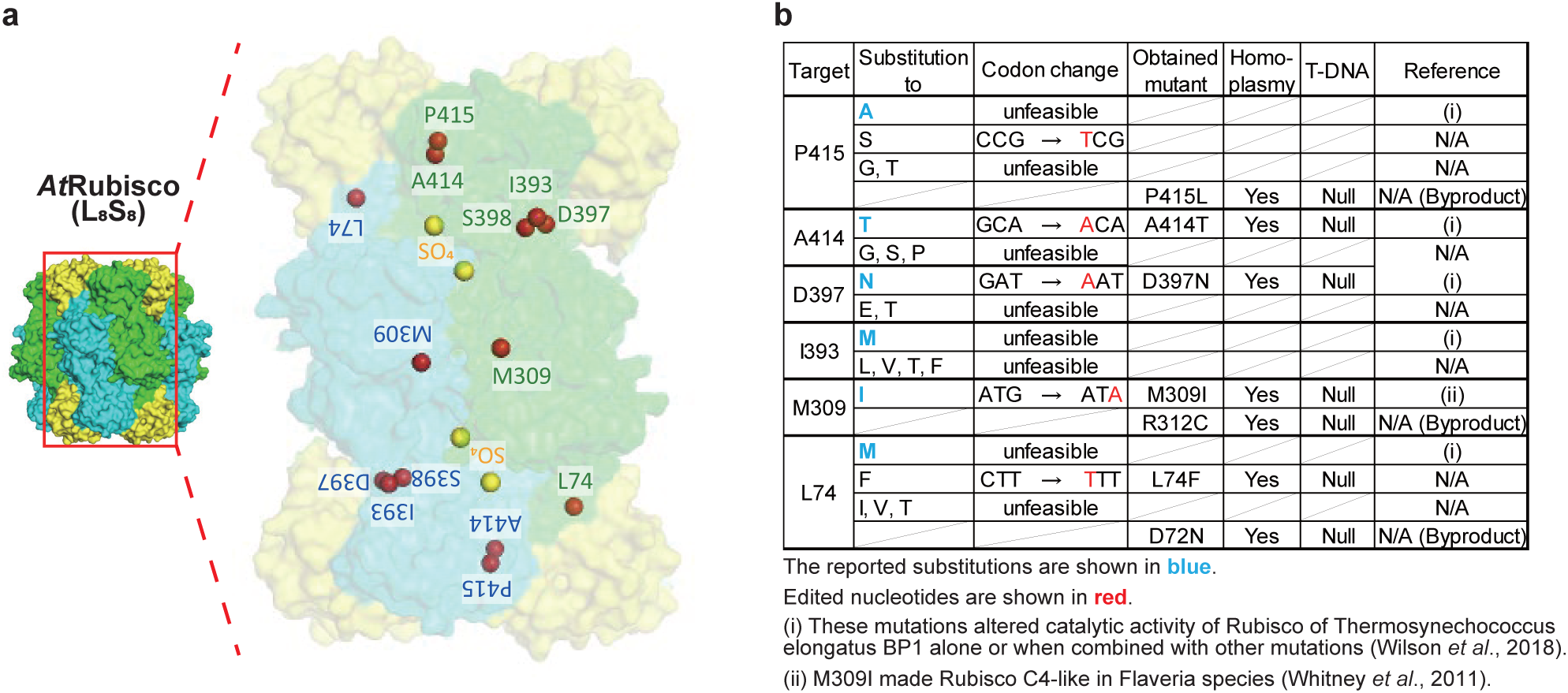
Amino acid substitutions in RbcL. **a**, Location of the target amino acids mapped onto the three-dimensional structure of *At*Rubisco. The RbcL dimer and four RbcS subunits are represented as semi-transparent surfaces and are colored in green, blue, and yellow, respectively. Sulfate molecules bound to the catalytic site are shown as yellow spheres to denote the catalytic site. The relative locations of the Cα atoms of the target amino acids are shown as red spheres. **b**, The target amino acids, desired substitutions, their causative codon changes, and obtained mutants and their genotypes. “unfeasible” means that this substitution is not achievable by C:G-to-T:A base editing. Homoplasmy was confirmed by genotyping both the T_2_ plants and their T_3_ progeny. Null indicates the absence of T-DNA.

The binary vectors encoding chloroplast base editors, ptpTALECD or ptpTALECD_v2mod [Fig. S1a], were integrated into the nuclear genome of *A. thaliana* Col-0. All the target substitutions seemed to be homoplasmically introduced (i.e., introduced across all chloroplast genomes, which are present in hundreds to thousands of copies per cell^22^) in one or more T_1_ plants (Fig. S1b-g and Table S2). In the T_2_ generation, only null-segregant plants, which lacked T-DNA were selected to avoid additional mutations induced by the base editors (Table S3). Including the unintended byproducts (D72N, R312C, and P415L mutants), a total of seven null-segregant single mutants were isolated (Table S3). All the T_3_ plants carried the homoplasmic mutation at the targets. Two T_3_ lines of M309I and D397N mutants were subjected to next-generation sequencing (NGS) to investigate off-target mutations in the chloroplast and mitochondrial genomes. There were no off-target mutations detected in three lines (Table S4), and a line rbcL_M309I-3_DddA11_11-1 had an off-target mutation of seemingly no effect, intergenic region, a C to T conversion located 65 bp upstream of the initiation codon of the *rpoB* gene. These results suggest that null-segregant lines with the homoplasmic mutation at the chloroplast targeted bases were generated. Subsequent analyses were conducted using T_4_ plants for the A414T mutants and T_3_ plants for the remaining mutants. For readability, the mutant lines will hereafter be referred to by shortened names, like “M309I Line 1” (listed in Table S5).

### Plant growth at ambient and high CO_2_ conc

A range of chloroplast-genome-edited *A. thaliana* plants targeting *rbcL* were cultivated at different CO_2_ concentrations including ambient concentration at 381 μmol mol^−1^ and high concentration at 549 μmol mol^−1^ (Fig. 2a, b). Both at the ambient and high CO_2_ concentration, M309I and D397N substitutions in *rbcL* exhibited significantly higher total leaf area and total shoot dry weight compared to Col-0 at 48 days after sowing (Fig. 2c-f), whereas at 24 or 25 days after sowing, they did not show any significant differences in total leaf area and total shoot dry weight (Fig. S2). All other lines exhibited either comparable or lower leaf area and dry weight relative to Col-0 irrespective of the growth CO_2_ concentration and timing after sowing (Figs. 2 and S2). Also, at ambient CO_2_ concentration, M309I and D397N exhibited significantly higher growth throughout a 90-day cultivation, with higher shoot dry weight, root dry weight, harvested seeds weight compared with Col-0 (Fig. S3, supplementary video).

**Fig. 2.**
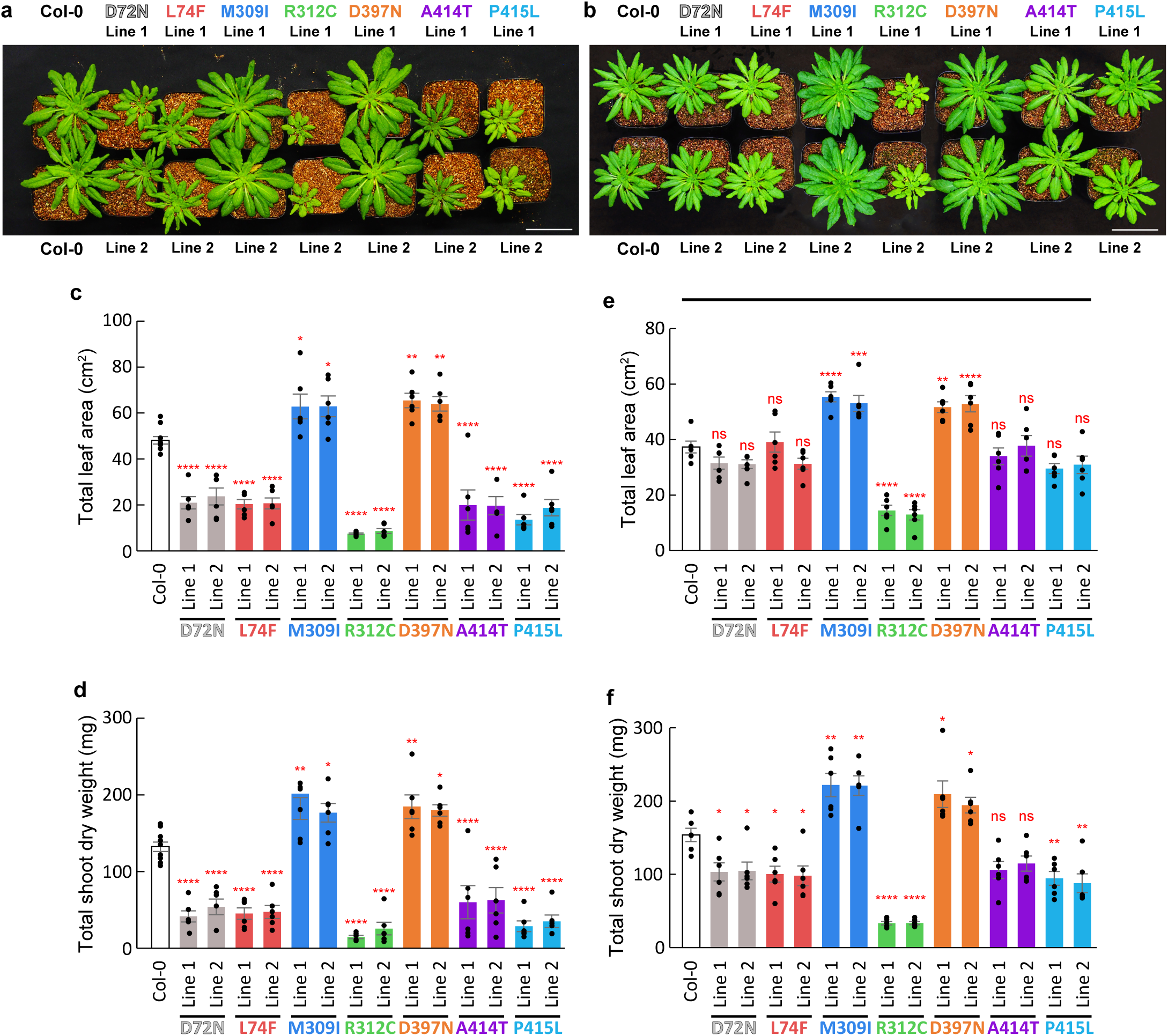
Plant growth at 48 days after sowing. a, **b**, The visual images of plants cultivated under ambient (381 µmol mol^-1^±14) (a) and high (549 µmol mol^-1^±23) (b) CO_2_ concentration. Bars = 5 cm. **c, d**, Total leaf area (c), and total shoot dry weight (d) of plants cultivated in ambient CO_2_ concentration. **e, f**, Total leaf area (e), and total shoot dry weight (f) of plants cultivated under high CO_2_ concentration. Data is mean ± SE. Individual data points were displayed as black dots. ns *p* > 0.05, * *p* < 0.05, ** *p* < 0.01, *** *p* < 0.001, **** *p* < 0.0001, n = 6 plants. Dunnett’s two-tailed test for multiple comparisons.

### Rubisco kinetics and photosynthetic components

M309I exhibited a significantly higher catalytic turnover rate (*k*_cat_) compared to Col-0 (Fig. 3a), but it also showed higher Michaelis-Menten constants for CO_2_ (*K*_c_), lower catalytic efficiencies for carboxylation (*k*_cat_/*K*_c_), and a lower ratio of carboxylation to oxygenation efficiencies (*S*_c/o_) (Fig. 3b-d). Moreover, D397N displayed a significantly higher *k*_cat_ and a lower *S*_c/o_ compared to Col-0 (Fig. 3a, d), while *K*_c_ and *k*_cat_/*K*_c_ were comparable (Fig. 3b, c). On the other hand, all other lines showed lower *k*_cat_ and *k*_cat_/*K*_c_ relative to Col-0 (Fig. 3a, c). Furthermore, for all lines, *K*_c_ was comparable to or higher than that of Col-0, whereas *S*_c/o_ was comparable to or higher than Col-0 (Fig. 3b, d).

**Fig. 3.**
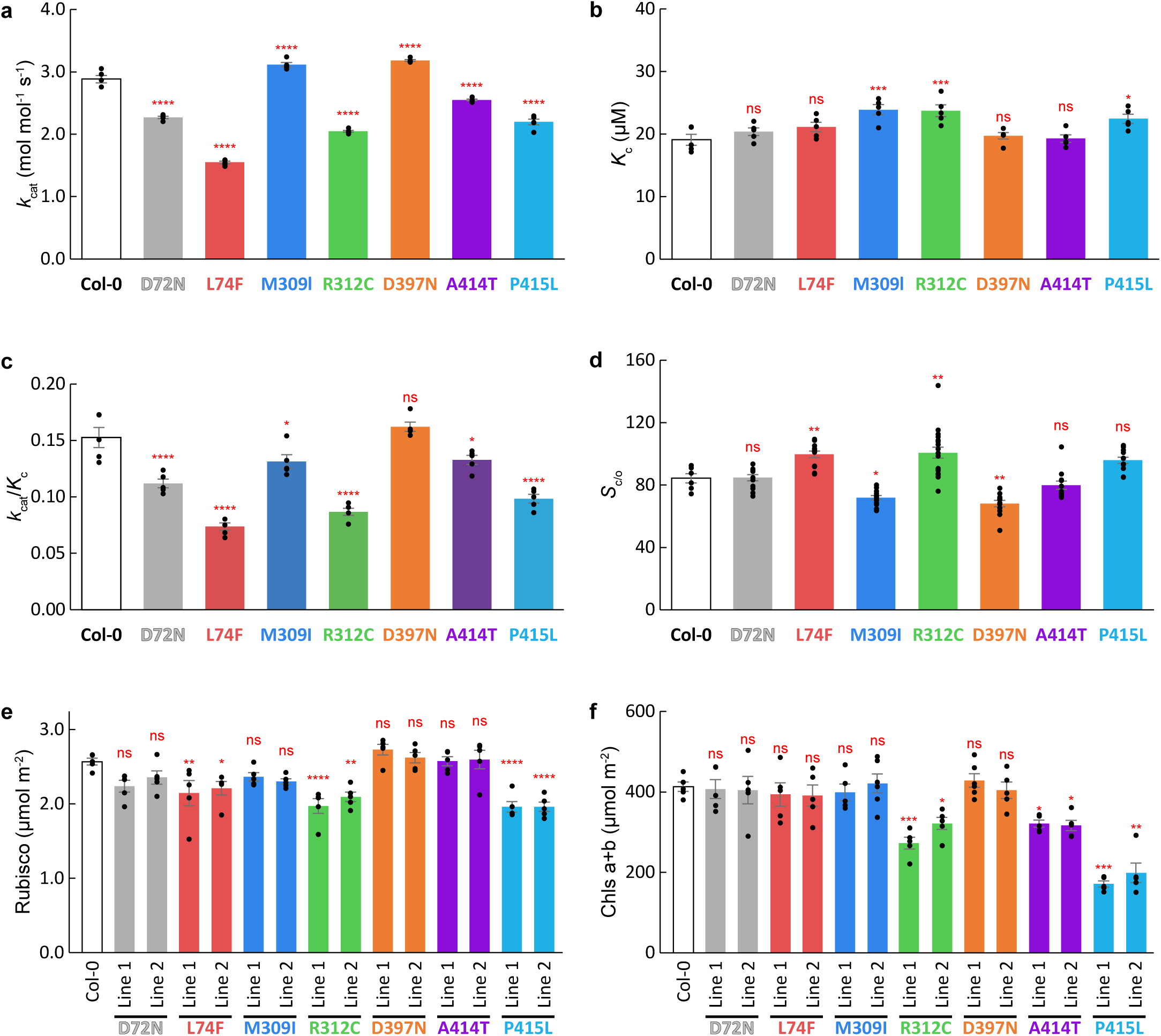
Rubisco kinetics, Rubisco content and Chlorophyll content in different lines. **a-d**, *k*_cat_ (a), *K*_c_ (b), *k*_cat_/*K*_c_ (c), and *S*_c/o_ (d) of Rubisco with various mutations. **e**, Rubisco content in different lines. **f**, Chlorophyll content in different lines. Data is mean ± SE. Individual data points were displayed as black dots. ns *p* > 0.05, * *p* < 0.05, ** *p* < 0.01, *** *p* < 0.001, **** *p* < 0.0001, n = 6 independent enzyme preparations. Dunnett’s two-tailed test for multiple comparisons.

D72N, M309I, D397N, and A414T exhibited Rubisco content comparable to that of Col-0, while other lines showed lower levels (Fig. 3e). Similarly, D72N, L74F, M309I, D397N, and A414T showed total chlorophyll (Chl) contents comparable to Col-0, whereas other lines had lower contents (Fig. 3f). The Chl a/b ratio was consistent across all lines except for P415L (Fig. S4). Also, no significant difference in Rubisco activase content was detected among Col-0, M309I, and D397N (Fig. S5).

### CO_2_ response of photosynthetic rate

The CO_2_ responses of several photosynthetic parameters were determined in all the chloroplast-genome-edited plants (Figs. 4a, b, and S6). At a CO_2_ concentration of 400 μmol mol^−1^, the M309I and D397N substitutions in *rbcL* exhibited a significantly higher CO_2_ assimilation rate and electron transport rate, and tended to have a lower NPQ compared to Col-0, whereas all other lines showed lower CO_2_ assimilation rates and electron transport rates relative to Col-0 (Fig. 4c-e). Furthermore, since stomatal conductance and intercellular CO_2_ concentrations for M309I and D397N were comparable to Col-0 (Fig. 4f, g), the intrinsic water-use efficiency (WUEi), calculated as the CO_2_ assimilation rate divided by stomatal conductance, was higher in M309I and D397N than in Col-0 (Fig. 4h). This finding indicates that overcoming photosynthetic limitations to improve Rubisco function resulted in increased water-use efficiency.

**Fig. 4.**
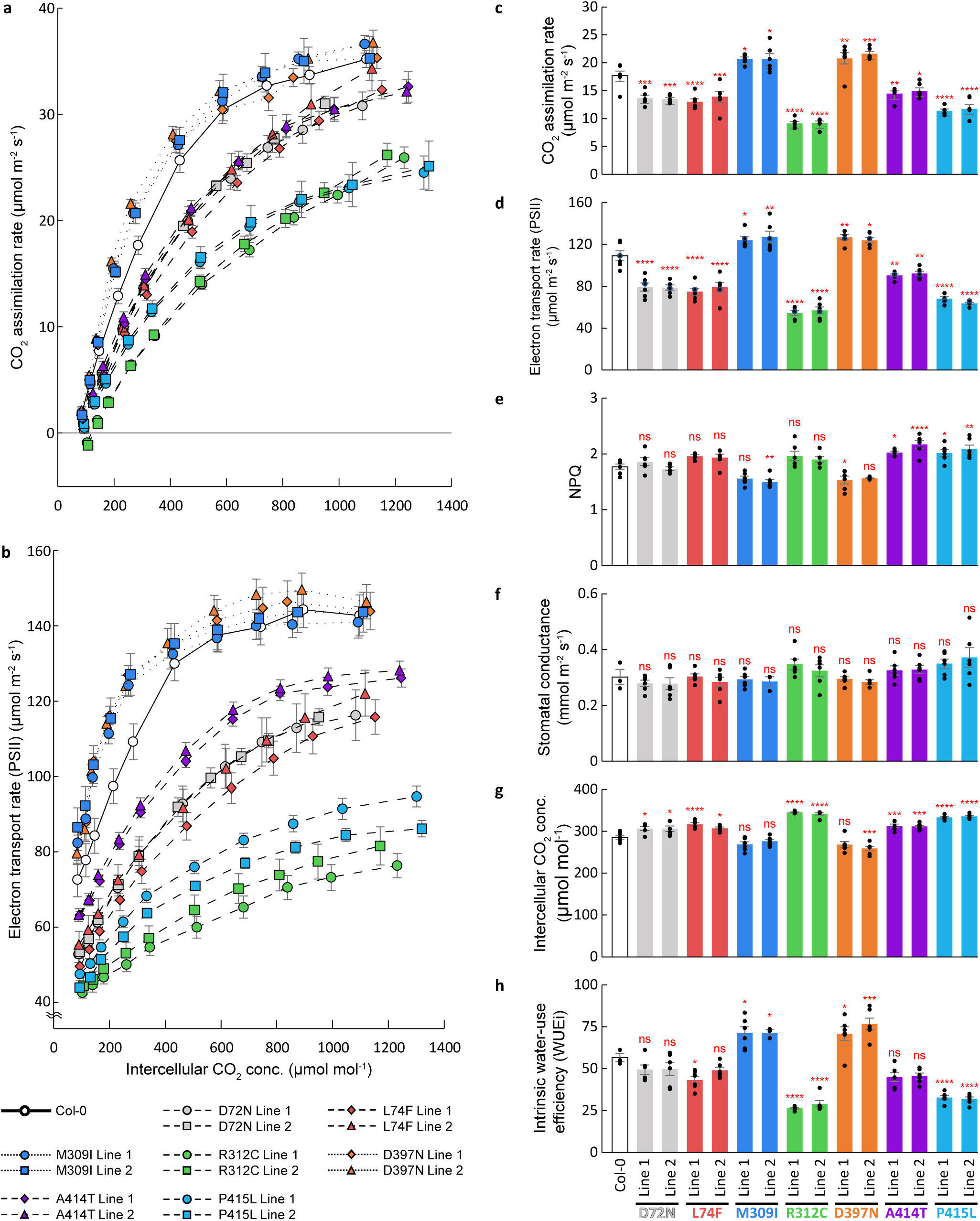
Photosynthesis parameters of different lines. **a**, Gas exchange parameters including CO_2_ assimilation rate of different lines in various intercellular CO_2_ concentrations. **b**, Photosynthetic fluorescence parameters including electron transport rate of Photosystem II (PSII) of different lines in various intercellular CO_2_ concentrations. **c-h**, Gas exchange and photosynthetic fluorescence parameters including CO_2_ assimilation rate (c), electron transport rate of PSII (d), NPQ (e), stomatal conductance rate (f), intercellular CO_2_ concentration (g), and intrinsic water-use efficiency (WUEi) (h) of different lines in 400 ppm CO_2_. Data is mean ± SE. Individual data points were displayed as black dots. ns *p* > 0.05, * *p* < 0.05, ** *p* < 0.01, *** *p* < 0.001, **** *p* < 0.0001, n = 6 plants. Dunnett’s two-tailed test for multiple comparisons.

On the other hand, at a CO_2_ concentration of 1200 μmol mol^−1^, the CO_2_ assimilation rate and electron transport rate were the same for all lines except R312C and P415L (Fig. S7a, b). Additionally, there were no significant differences in photosynthetic parameters, including stomatal conductance, intercellular CO_2_ concentration, WUEi, and NPQ, across all lines (Fig. S7c-f).

### Structures of M309I and D397N Rubisco

To address how M309I and D397N mutations of RbcL affect the Rubisco activity, we purified Col-0, M309I, and D397N Rubisco, and collected cryo-EM data in the presence of 50 mM sulfate ions, in analogy with the previous crystallographic study^14^. Rubisco structure is related to D4 symmetry, and the cryo-EM density maps imposed with D4 symmetry exhibited the highest resolution. Although we also fit the models into EM density without imposing symmetry (Extended Data Table 1-5), no substantial differences were observed compared to the symmetric models. Therefore, the discussion hereafter will focus on the models imposed with D4 symmetry. The Col-0, M309I, and D397N Rubisco models were determined at resolutions of 1.76, 1.75, and 1.74 Å, respectively (Fig. 5, and Extended Data Table 1 and summarized in Extended Data Figs. 1-3).

**Fig. 5.**
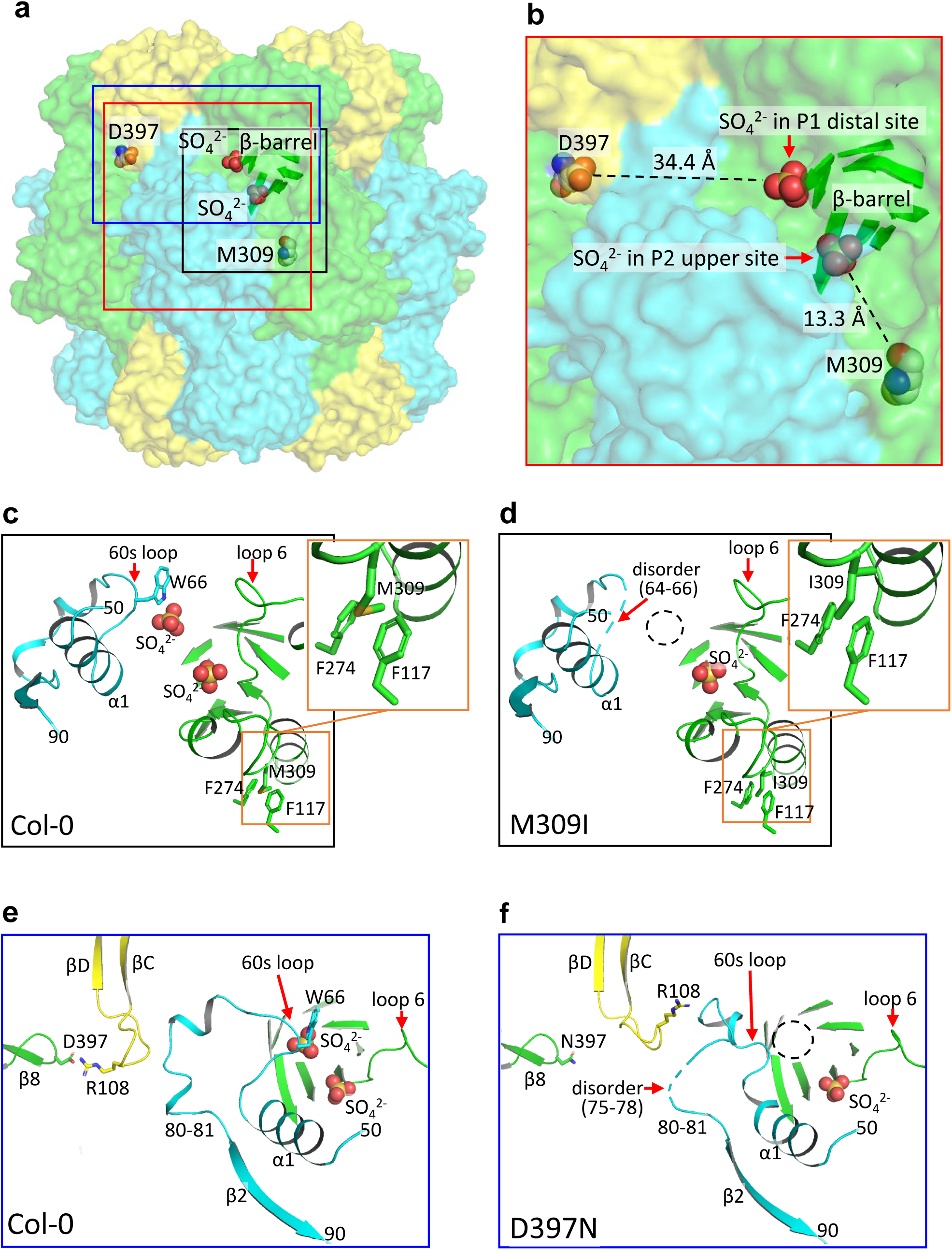
*A. thaliana* Rubisco structures in complex with sulfate ions. a, Location of M309 and D397 in Col-0 Rubisco. Eight RbcL are colored green and cyan, and eight RbcS are colored yellow. M309, D397, and two sulfate ions are drawn in space-filling mode. The β-barrel whose C-terminal regions form the active site is shown. The red, black, and blue squares indicate the focused area of **b**, **c** and **d**, and **e** and **f**, respectively. **b,** Focused structure around the active site. Distances between M309 and SO_4_^2-^ in the P2 upper site and D397 and SO_4_^2-^ in the P1 distal site are shown. **c, d,** Structural comparison of the active site between Col-0 and M309I. Key interacting residues are labeled. The dashed circle highlights the P1 distal site, where no sulfate ion is present. **e, f,** Structural comparison of the active site between Col-0 and D397N.

Expressed in leaves of *A. thaliana* are four distinct isoforms of RbcS (RBCS1A, RBCS1B, RBCS2B, and RBCS3B), which possibly correlate with the accumulation of Rubisco in chloroplasts within the leaves^23^ (Extended Data Fig. 4). Growth temperature affects the expression levels of these RbcS isoforms with distinct kinetics^24^. Given our growth conditions (see Methods), RBCS1A is expected to be the most abundant, and RBCS3B is the second^23^. The cryo-EM density maps represent a mixture of these isoforms of RbcS (Extended Data Fig. 5). Our SDS-PAGE analysis also shows that the RBCS1A level accounted for approximately half of the total (Fig. S8), consistent with the previous report^23^. As RBCS1A is deduced to be the most abundant, we decided to fit *A. thaliana* RbcS structures based on the amino acid sequence of RBCS1A.

The folding of M309I and D397N Rubisco are basically the same as those of Col-0 Rubisco (Extended Data Fig. 6). The Col-0, M309I, and D397N Rubisco structures contain sulfate ions at the RuBP phosphate group binding sites, due to the presence of 50 mM sulfate ions in the cryo-EM specimen solutions. In the Col-0 Rubisco active site, two sulfate ions were observed at the P1 distal site and P2 upper site (Fig. 5c), whereas in M309I and D397N, only one sulfate ion was present in the P2 upper site (Figs. 5d, f, and S9). In Col-0 Rubisco, the side chain of W66 interacts with the sulfate at the P1 distal site, but such interactions are not observed in M309I and D397N.

The two catalytic loops of Rubisco, namely 60s loop (residues 64-67) and loop 6 (residues 332-337), typically undergo a transition from disorder to an ordered state upon binding of the sugar-phosphate substrate [or the transition state analogue 2-carboxy-D-arabinitol-1,5-bisphosphate (2CABP)]^25^. In our condition with 50 mM sulfate ions, all of Rubisco’s loop 6 were structurally ordered with an open conformation (Fig. 5c-f). In contrast, Col-0 and D397N Rubisco’s 60s loop were structurally ordered but was disordered in M309I (Fig. S10). Notably, D397N has a 60s loop with distinct conformation (Figs. 5f, and S9b).

## Discussion

The transformation of chloroplast genomes in plants offers a promising approach to enhancing photosynthesis and crop productivity, as many chloroplast genome-encoded genes are involved in photosynthesis. Despite extensive research over the past fifty years aimed at improving Rubisco’s catalytic properties, efforts to engineer vascular plants by replacing them with “super Rubisco” enzymes have largely been unsuccessful and often resulted in reduced plant growth. This is primarily due to challenges in coordinating enzyme expression within its intricate plant metabolic and protein structure relationships. This study demonstrates that Rubisco kinetics can be significantly enhanced through the least change, single amino acid substitutions at M309I and D397N, each of which increases *k*_cat_ of Rubisco (Fig. 3). M309 and D397 are located at the RbcL-RbcL and RbcL-RbcS interfaces, respectively, both of which are distant from the active site (Fig. 5b). The molecular mechanisms by which these mutations at sites distant from the active site affect Rubisco activity have remained unclear so far^26^. In the present structural analysis, we observed structural differences in the 60s loop between the Col-0, M309I, and D397N. Interestingly, the structural changes of the 60s loop have also been implicated in hybrid Rubisco with the replacement of rice Rubisco RbcS by sorghum counterparts with increased *k*_cat_^14^. Thus, mutations in amino acids that influence the structure of the 60s loop may be the key to enhancing Rubisco activity. However, predicting and designing mutations to achieve such structural alternations remains challenging (see Supplementary Discussion for a more detailed discussion).

The structural changes caused by the M309I and D397N substitutions improved Rubisco kinetics (Fig. 3), and the engineering Rubisco with a high carboxylation rate into plant chloroplasts presents a viable strategy for improving photosynthesis (Fig. 4c), WUEi (Fig. 4h), and plant growth (Fig. 2) under both ambient and elevated CO_2_ concentrations. Rubisco has evolved to enhance *S*_c/o_, *k*_cat_, and *k*_cat_/*K*_c_ in angiosperms. However, the evolution rate of the *rbcL* gene is known significantly slower compared to other gene/protein sequences^27^. Naturally, this slow rate of gene evolution has resulted in Rubisco’s characteristics not keeping pace with the increased atmospheric CO_2_ concentrations since the Industrial Revolution. In this study, we utilized the ptpTALECD genome editing tool^21^ for chloroplast genome editing to accelerate Rubisco evolution, introducing properties adapted to high CO_2_ environments, similar to C4 Rubisco. M309 and D397 are well conserved in diverse plant species even at the nucleotide sequence level (Table S1). Consequently, the introduction of M309I and D397N mutations may improve biomass production in most herbaceous and woody plants, including agriculturally important crops. This approach, not a drastic change but the least change of base editing, led to improvements in photosynthesis and plant growth (Figs. 2, 4). Many of the previous approaches aim to overcome Rubisco’s limitations through various bioengineering strategies, either by enhancing chloroplast CO_2_ levels^28,29^ or by improving Rubisco’s catalytic properties (this study and two previous studies^3,30^). As, in recent years, CO_2_ levels have been rising more rapidly than humanity had anticipated, Rubisco modification itself, even without concentrating CO_2_ within the chloroplasts, may be able to boost photosynthetic capacity some more. Chloroplast transformation offers several advantages, including precise genome manipulation via homologous recombination, avoidance of transgene silencing, and better control over the escape of transgenes into the environment^31^. As chloroplast genome editing can be performed in plant species that allow nuclear genome-targeted transformation, this transformation strategy could also be applied to introduce point mutations in other chloroplast-encoded genes in a non-GMO context in many countries, potentially driving an acceptable and feasible new green revolution.

## Methods

### Sequence analysis of RbcL

*rbcL* sequences were extracted from reference genomes (Table S1) and compared using Geneious Prime v2024.0.7.

### Development and Introduction of Rubisco Mutants

The sequences that the TALE arrays bind to (listed in Table S6) were settled based on the positional tendencies of edited cytosines^21,32^. All of the mutants except for M309I were obtained by ptpTALECD. The M309I mutation could not be achieved with ptpTALECD; therefore, ptpTALECD_v2mod, which has a cytidine deaminase domain with a higher base-editing activity than that of ptpTALECD was used to introduce this mutation. Binary vectors to express ptpTALECD and ptpTALECD_v2mod were assembled as described previously^21,32^, but the vector of ptpTALECD_v2mod was assembled using intermediate plasmids (Addgene ID #191599-191602, and 191607- 191610) instead of those employed to assembly ptpTALECD vectors (Addgene ID #171724-171735, 191579-191598). The binary vectors were integrated into the nuclear genome of *A. thaliana* Col-0 by floral dipping^33^. T_1_ seeds were selected based on the presence of GFP fluorescence^34^. DNA was roughly extracted from a T_1_ leaf by incubating it with 50 µL of TE Buffer [100 mM Tris-HCl (pH 9.5) and 10 mM ethylenediaminetetraacetic acid (EDTA) (pH 8.0) in diluted water] at 98℃ for 15 minutes. Targeted sequences were amplified from the DNA using KOD one PCR Master Mix (TOYOBO). PCR products were purified with FastGene Gel/PCR Extraction Kit (Nippon Genetics) and subsequently subjected to Sanger sequencing. PCR primers are listed in Table S7. Null-segregant T_2_ seeds were selected based on the absence of GFP fluorescence. Genotyping of T_2_ plants was conducted using the same methods as for the T_1_ plants.

### NGS analyses

To assess off-target mutations in chloroplast and mitochondrial genomes, NGS analyses were performed by using total DNAs extracted from a leaf of T_3_ null-segregant plants with Maxwell RSC Plant DNA Kit (Promega). Siblings of these plants were subjected to the phenotypic assays in this study. We ordered Macrogen Japan to prepare paired-end libraries using a Nextera Flex DNA library Prep Kit (Illumina) and to sequence the libraries using Illumina NovaSeq X platform. NGS genome data of Col-0 was obtained from our previous study^32^. The subsequent processes to analyze the NGS data, including trimming of low-quality and adaptor sequences in the reads, mapping of the reads to the reference chloroplast and mitochondrial genome sequences (AP000423.1 and BK010421.1, respectively), and SNP calling, were conducted using Geneious Prime v2024.0.7. We filtered out inadequately mapped reads with mapping identities ≤97% or alignment cover rates ≤80%. Finally, we listed base positions where the difference between the substitution rate of a T_3_ sample and the average substitution rate of the two Col-0 samples was ≥5%.

### Plant material and cultivation conditions

To establish genome-edited lines, seedlings of *A. thaliana* ecotype Col-0, and T_2_ and T_3_ plants were grown on 1/2 Murashige-Skoog (MS) agar plates for 2-3 weeks before being transferred to Jiffy-7 (Jiffy Products International). The plants were grown under 22℃ under long-day conditions (a daylight/dark period of 16/8 h). T_1_ seeds were sown on 1/2 MS agar plates containing 125 mg L^-1^ Claforan, and T_1_ plants were grown under similar conditions. The composition of the 1/2 MS plates was described previously^35^. For the phenotyping assays, the seeds of Arabidopsis were sown in a 1:1 mixture of MetroMix (Metromix 350, Hyponex) and vermiculite, and cultivated in several environment-controlled growth chambers. The growth chambers were operated with a daylight/dark period of 10/14 h, and photosynthetic photon flux density (PPFD) in the daylight period was set to 150 μmol photons m^-2^ s^-1^. Air temperature and relative humidity were set to a constant 22°C and 60%, respectively. Chamber CO_2_ concentrations were set to ambient concentration (381 ± 14 μmol mol^−1^) and high concentration (549 ± 23 μmol mol^−1^).

### Plant growth

Plants were cultivated under two different CO_2_ conditions: ambient CO_2_ concentration and high CO_2_ concentration. For early-stage growth determinations, plants were harvested after 25 days at both ambient and high CO_2_ concentrations. For later-stage growth determinations, plants were harvested after 48 days at both ambient and high CO_2_ concentrations. During harvest, all above-ground tissues were separated, and the leaves were scanned using a scanner. Leaf area was measured using ImageJ software. After scanning, all above-ground tissues were collected and dried at 60°C for 48 hours to determine their dry weight. Below-ground tissues were carefully uprooted without damaging the root systems, rinsed in running tap water, and gently rubbed to remove any soil residues. Cleaned root tissues were then dried at 60°C for 48 hours, and their dry weight was measured.

### Photosynthetic responses to CO_2_ concentrations

The gas exchange rate and chlorophyll fluorescence were measured simultaneously in fully expanded young leaves attached to 6- to 8-week-old plants grown at ambient CO_2_ concentration with a portable gas exchange system (LI-6400XT, LI-6400-40 leaf chamber fluorometer, LI-COR). The CO_2_ assimilation rate, stomatal conductance, and intercellular CO_2_ concentration, together with chlorophyll fluorescence parameters including electron transport rate at Photosystem II and non-photochemical quenching (NPQ), were measured at a PPFD of 1000 μmol photons m^−2^ s^−1^, with relative humidity of 60-70%, temperature of 25°C, and CO_2_ concentrations of 50 μmol mol^−1^ to 1500 μmol mol^−1^. To determine Rubisco CO_2_/O_2_ specificity (*S*_c/o_), the gas exchange rates were measured under the conditions as described above with two different light intensities at a PPFD of 125 and 500 μmol photons m^−2^ s^−1^. The *S*_c/o_ was calculated using the CO_2_ compensation points independent of photorespiration estimated by the intersection of these two regression lines^36^.

### Rubisco, Rubisco activase, and photosynthetic pigments

Leaf tissues from the same leaf used for gas-exchange measurement were sampled immediately after gas exchange measurements, frozen in liquid nitrogen, and stored at −80°C until analysis. The Rubisco content was determined by formamide extraction of SDS-PAGE-separated, Coomassie Brilliant Blue R-250–stained bands corresponding to the large and small subunits of Rubisco, with Bovine Serum Albumin (BSA) as the standard^37^. The Rubisco activase content was quantified by immunoblotting of SDS-PAGE-separated proteins with Primary anti-activase antibody, and secondary anti-rabbit IgG HRP-linked antibody (GE Healthcare, NA934), on a polyvinylidene difluoride membrane^38^. Chlorophyll was extracted in 80% v/v acetone, and chlorophyll content was determined as described previously^39^.

### Rubisco kinetics

The leaves were homogenized in extraction buffer containing 100 mM Bicine-NaOH, 1 mM EDTA, 5 mM MgCl_2_, 2 mM NaH_2_PO_4_, 5 mM dithiothreitol (DTT), 20 mM ascorbate, 4 mM amino-n-caproic acid, 0.8 mM benzamidine, 0.4% (w/v) bovine serum albumin, and 1% (w/v) polyvinylpolypyrrolidone, pH 8.0, using a chilled mortar and pestle. The homogenate was then centrifuged at 15,000 × g for 2 min at 4°C. Rubisco in the supernatant was activated by pre-incubation with 15 mM MgCl_2_ and 5 mM NaHCO_3_ on ice for 15 min. Rubisco activity was determined at 25°C using [^14^C] NaHCO_3_ (specific activity, 37 MBq mmol^-1^) by assaying the incorporation of ^14^C into acid-stable products, as described previously^14^. The reaction was started by the addition of activated Rubisco to a reaction mixture containing 100 mM Bicine-NaOH, 20 mM MgCl_2_, 1 mM EDTA, 5 mM DTT, 0.5 mM RuBP, 1.0 W-A units carbonic anhydrase, and 0.5-20 mM NaH^14^CO_3_, pH 8.2. After 1 min, 1/2 volume of 1 N HCl was added to the reaction solution to stop the reaction. The acidified reaction mixes were dried and acid-stable ^14^C was measured by liquid scintillation. The Rubisco catalytic site concentration was determined by measuring the stoichiometric binding of [^14^C] 2- carboxyarabinitol bisphosphate (specific activity, 1.85 GBq mmol^-1^) as described previously^40^. The *k*_cat_ and *K*_c_ were calculated by direct fitting of the Rubisco activities at six different NaH^14^CO_3_ concentrations (0.5-20 mM) to the Michaelis-Menten equation using the KaleidaGraph data analysis software (Synergy Software).

### Protein purification, cryo-EM specimen preparation, and data collection

Col-0, M309I, and D397N Rubisco were purified from leaves in Col-0 and chloroplast-genome-edited *A. thaliana* as described in the previous studies for rice Rubisco^41^, with the following modifications. Rubisco fractions were applied twice to a HiLoad 16/60 Superdex 200 prep grade column (Cytiva). In the final step of the gel filtration, we used buffer containing 80 mM Hepes-KOH (pH 8.0 at 20°C), 1 mM EDTA, and 5 mM DTT. The purified enzyme was concentrated by centrifugation (Vivaspin 20, MWCO: 50,000, Sartorius) up to 15 mg mL^-1^ for Rubisco, frozen with liquid N2, and stored at-80 °C until the cryo-EM experiments. The amount of Rubisco was estimated spectrophotometrically by assuming an extinction coefficient of 1.67 absorbance units for 1 mg mL^-1^ at 280 nm.

A 3 µL aliquot of Rubisco (1 mg mL^-1^) in the buffer containing 80 mM Hepes-KOH pH 8.0, 5 mM DTT, 1 mM EDTA, and 50 mM ammonium sulfate was applied to Quantifoil Cu R1.2/1.3 holey carbon grids and frozen in liquid ethane using a Vitrobot IV system (Thermo Fisher Scientific, 4°C and 100 % humidity, 3 s blotting time). The grids were inserted into a CRYO ARM 300 transmission electron microscopy (JEOL Ltd. Japan) equipped with a cold field-emission electron gun operated at 300 kV and an Ω-type energy filter with a 20-eV slit width. Cryo-EM images were recorded with a K3 direct electron detector camera (Gatan, USA) at a nominal magnification of ×60,000, corresponding to an image pixel size of 0.87 Å with a setting defocus range from-0.7 to-2.2 μm, using Serial-EM^42^. The holes were detected using YoneoLocr^43^. Movie frames were recorded in CDS counting mode with a total exposure of 3 s and a total dose of ∼80 electrons Å^-2^. Each movie was fractionated into 40 frames. A total number of 2,732, 3,258 and 3,509 movies were collected for Col-0, M309I and D397N of the Rubisco, respectively.

### Cryo-EM image processing and model building

The Process of single particle analysis of the Col-0 Rubisco was performed using RELION 4.0^44^. Image processing procedures of Col-0 Rubisco are described in Extended Data Fig. 1. After performed motion correction to align all micrographs, followed by the estimation of parameters of the contrast transfer fraction (CTF), the selected particles were extracted into a box of 80 × 80 pixels with binning ×4. The coordinate data used for particle picking (1,487,277) was obtained by BoxNet’s particle picking of Warp^45^ performed during data collection. 1,214,112 particle images from good 2D class average images were selected for the initial 3D model with C1 symmetry into three classes. After initial 3D model, a good initial volume classes were re-extracted into a box of 320 × 320 pixels (unbinned). After 3D refinement with C1 symmetry, CTF refinement and Polish process, particle images were performed 3D refinement with D4 symmetry. 737,173 particles were subjected to 3D classification with C1 symmetry into three classes without alignment. Following 3D refinement with D4 symmetry, the map resolution reached 1.76 Å [the Fourier shell correlation (FSC) = 0.143]. After 3D refinement and CTF refinement for each of these classes, postprocessing yielded the 3D maps with resolutions of 2.42 Å (89,314 particles), 2.02 Å (274,725 particles) and 2.08 Å (353,069 particles), respectively, according to 0.143 criterion of FSC.

The process of single particle analysis of the M309I and D397N Rubisco was also performed using RELION 4.0 in the same way as for Col-0 Rubisco (Extended Data Figs. 2 and 3, respectively)^44^. Particle picking (2,074,623, 2,302,408) was initially obtained by BoxNet’s particle picking of Warp^45^, respectively. In M309I, following 3D refinement with D4 symmetry, the map resolution reached 1.75 Å (FSC = 0.143). After 3D refinement and CTF refinement for each three classes, postprocessing yielded the 3D maps with resolutions of 2.01 Å (338,656 particles), 1.95 Å (578,985 particles), and 2.04 Å (252,690 particles), respectively, according to 0.143 criterion of FSC. In D397N, following 3D refinement with D4 symmetry, the map resolution reached 1.74 Å (FSC = 0.143). After 3D refinement and CTF refinement for each three classes, postprocessing yielded the 3D maps with resolutions of 2.10 Å (258,006 particles), 2.01 Å (306,772 particles), and 2.01 Å (339,707 particles), respectively, according to 0.143 criterion of FSC.

The models were built using the crystal structure of sulfate-bound rice Rubisco (PDB code: 6KYI^14^) as initial models. After the initial models were manually fitted into the map using UCSF Chimera ver. 1.16^46^ and modified in Coot ver. 0.9.6^47^, real-space refinement was performed in PHENIX ver. 1.20.1^48^. The model was validated using MolProbity ver. 4.5.2^49^ in PHENIX ver. 1.20. 1^48^, and this cycle was repeated several times. Cryo-EM data collection, refinement, and validation statistics are summarized in Extended Data Table 1-5.

The stained bands were scanned and processed using ImageJ ver. 1.53q^50^. Figures were prepared using UCSF Chimera ver. 1.16^46^, CLUSTALX ver. 2.0^51^, ESPript ver.

3.0^52^ (https://espript.ibcp.fr), and PyMOL ver. 2.5.0 (Schrödinger, LLC, USA).

## Statistical analysis

The significance of variations between means of multiple lines and the control group was evaluated by Dunnett’s two-tailed test. Statistical analysis was performed using Prism v. 8.0.1 software.

## Data availability

Cryo-EM atomic coordinates and maps of Col-0, M309I, and D397N Rubisco have been deposited in the Protein Data Bank (PDB) and the Electron Microscopy Data Bank (EMDB) under the accession codes 9L58 and EMD-62824, 9L5D and EMD-62829, and 9L5I and EMD-62834 for three classes merged with D4 symmetry; 9L57 and EMD-62823, 9L5D and EMD-62829, and 9L5H and EMD-62833 for three classes merged with C1 symmetry; 9L59 and EMD-62825, 9L5E and EMD-62830, and 9L5J and EMD-62835 for class1 with C1 symmetry; 9L5A and EMD-62826, 9L5F and EMD-62831, and 9L5K and EMD-62836 for class2 with C1 symmetry; 9L5B and EMD-62827, 9L5G and EMD-62832, and 9L5L and EMD-62837 for class3 with C1 symmetry, respectively. The other coordinates used in this study are available from PDB. Source data are provided in this paper.

## Acknowledgements

This work was supported by: JSPS KAKENHI grant 21H02171 (W.Y.), 24H02271 (S.A.), 24H02277 (W.Y., H.F., H.M.), 24K01994 (H.M.), 23K18033(H.M.) JPJSCCA20230008 (S.A.), 22J20237(I. N.); JST OPERA (Open Innovation with Enterprises, Research Institute and Academia) grant JPMJOP1861 (K.N.); the Program for the R-GIRO Research from the Ritsumeikan Global Innovation Research Organization, Ritsumeikan University (H.M.); AMED BINDS (Platform Project for Supporting Drug Discovery and Life Science Research (BINDS) grant JP21am0101117 and JP22ama121003 (K.N.), and JP23ama121001 (H.M.); AMED CiCLE (Cyclic Innovation for Clinical Empowerment) grant JP17pc0101020 (K.N.); JEOL YOKOGUSHI Research Alliance Laboratories of Osaka University (K.N.); the Cooperative Research Program of the Institute for Protein Research, Osaka University (CR-22-02 and CR-23-02). We thank Yoshiko Tamura and Reiko Masuda for their genome editing technical support.

## Author contributions

W.Y, I.N., H.M and S.A. conceived and designed the study. I.N. and S.A. constructed the vectors, established chloroplast genome-edited lines, and carried out related DNA analyses. W.Y., Q.Y., Y.S., and H.F. performed analysis of *in vivo* Rubisco assay, photosynthesis and plant growth, and F. H. performed *in vitro* Rubisco assay. R.U., Y.N., and W.Y. prepared the protein sample, and T.M., R.U., Y.N., and K.N. performed data collection, processing and H.M performed model building and refinement. W.Y, I.N., H.F. H.M and S.A. wrote the manuscript. All authors discussed the results and revised the manuscript.

## Additional information

Correspondence and requests for materials should be addressed to, and (W. Y.; 0000-0001-7215-4736, H. M.; 0000-0003-0361-3796 and S. A.; 0000-0002-9537-1626).

## References

1. Anwar, M. R., Liu, D. L., Macadam, I. & Kelly, G. Adapting agriculture to climate change: a review. Theor Appl Climatol 113, 225–245 (2013).

2. Pachauri, R. K. et al. Climate Change 2014: Synthesis Report. Contribution of Working Groups I, II and III to the Fifth Assessment Report of the Intergovernmental Panel on Climate Change. (Ipcc, 2014).

3. Evans, J. R. Improving photosynthesis. Plant physiology 162, 1780–1793 (2013).

4. Long, S. P., Marshall-Colon, A. & Zhu, X.-G. Meeting the global food demand of the future by engineering crop photosynthesis and yield potential. Cell 161, 56–66 (2015).

5. Zhu, X.-G., Long, S. P. & Ort, D. R. Improving Photosynthetic Efficiency for Greater Yield. Annu. Rev. Plant Biol. 61, 235–261 (2010).

6. Long, B. M. et al. Carboxysome encapsulation of the CO2-fixing enzyme Rubisco in tobacco chloroplasts. Nature communications 9, 1–14 (2018).

7. Zhu, X.-G., Portis, A. R.&Long, S. P. Would transformation of C_3_ crop plants with foreign Rubisco increase productivity? A computational analysis extrapolating from kinetic properties to canopy photosynthesis. Plant Cell & Environment 27, 155–165 (2004).

8. Sharwood, R. E., Ghannoum, O. & Whitney, S. M. Prospects for improving CO2 fixation in C3- crops through understanding C4-Rubisco biogenesis and catalytic diversity. Current opinion in plant biology 31, 135–142 (2016).

9. Sharwood, R. E. Engineering chloroplasts to improve Rubisco catalysis: prospects for translating improvements into food and fiber crops. New Phytologist 213, 494–510 (2017).

10. Martin-Avila, E. et al. Modifying plant photosynthesis and growth via simultaneous chloroplast transformation of Rubisco large and small subunits. Plant Cell 32, 2898–2916 (2020).

11. Conlan, B. & Whitney, S. Preparing Rubisco for a tune up. Nature plants 4, 12–13 (2018).

12. Flamholz, A. I. et al. Revisiting Trade-offs between Rubisco Kinetic Parameters. Biochemistry 58, 3365–3376 (2019).

13. Davidi, D. et al. Highly active rubiscos discovered by systematic interrogation of natural sequence diversity. The EMBO Journal 39, e104081 (2020).

14. Matsumura, H. et al. Hybrid Rubisco with complete replacement of rice Rubisco small subunits by sorghum counterparts confers C4 plant-like high catalytic activity. Molecular Plant 13, 1570– 1581 (2020).

15. Lin, M. T., Occhialini, A., Andralojc, P. J., Parry, M. A. & Hanson, M. R. A faster Rubisco with potential to increase photosynthesis in crops. Nature 513, 547–550 (2014).

16. Occhialini, A., Lin, M. T., Andralojc, P. J., Hanson, M. R. & Parry, M. A. J. Transgenic tobacco plants with improved cyanobacterial Rubisco expression but no extra assembly factors grow at near wild-type rates if provided with elevated CO_2_. The Plant Journal 85, 148–160 (2016).

17. Gunn, L. H., Martin Avila, E., Birch, R. & Whitney, S. M. The dependency of red Rubisco on its cognate activase for enhancing plant photosynthesis and growth. Proc. Natl. Acad. Sci. U.S.A. 117, 25890–25896 (2020).

18. Orr, D. J. et al. Hybrid cyanobacterial-tobacco Rubisco supports autotrophic growth and procarboxysomal aggregation. Plant Physiology 182, 807–818 (2020).

19. Wilson, R. H., Martin-Avila, E., Conlan, C. & Whitney, S. M. An improved Escherichia coli screen for Rubisco identifies a protein–protein interface that can enhance CO2-fixation kinetics. Journal of Biological Chemistry 293, 18–27 (2018).

20. Whitney, S. M. et al. Isoleucine 309 acts as a C4 catalytic switch that increases ribulose-1, 5- bisphosphate carboxylase/oxygenase (rubisco) carboxylation rate in Flaveria. Proceedings of the National Academy of Sciences 108, 14688–14693 (2011).

21. Nakazato, I. et al. Targeted base editing in the plastid genome of Arabidopsis thaliana. Nature Plants 7, 906–913 (2021).

22. Zoschke, R., Liere, K. & Börner, T. From seedling to mature plant: Arabidopsis plastidial genome copy number, RNA accumulation and transcription are differentially regulated during leaf development. The Plant Journal 50, 710–722 (2007).

23. Izumi, M., Tsunoda, H., Suzuki, Y., Makino, A. & Ishida, H. RBCS1A and RBCS3B, two major members within the Arabidopsis RBCS multigene family, function to yield sufficient Rubisco content for leaf photosynthetic capacity. Journal of Experimental Botany 63, 2159–2170 (2012).

24. Cavanagh, A. P., Slattery, R. & Kubien, D. S. Temperature-induced changes in Arabidopsis Rubisco activity and isoform expression. Journal of experimental botany 74, 651–663 (2023).

25. Duff, A. P., Andrews, T. J. & Curmi, P. M. The transition between the open and closed states of rubisco is triggered by the inter-phosphate distance of the bound bisphosphate. Journal of Molecular Biology 298, 903–916 (2000).

26. Andersson, I. & Backlund, A. Structure and function of Rubisco. Plant Physiology and Biochemistry 46, 275–291 (2008).

27. Bouvier, J. W., Emms, D. M. & Kelly, S. Rubisco is evolving for improved catalytic efficiency and CO_2_ assimilation in plants. Proc. Natl. Acad. Sci. U.S.A. 121, e2321050121 (2024).

28. Peterhansel, C., Blume, C. & Offermann, S. Photorespiratory bypasses: how can they work? Journal of Experimental Botany 64, 709–715 (2013).

29. Price, G. D. et al. The cyanobacterial CCM as a source of genes for improving photosynthetic CO2 fixation in crop species. Journal of experimental botany 64, 753–768 (2013).

30. Parry, M. A. et al. Rubisco activity and regulation as targets for crop improvement. Journal of experimental botany 64, 717–730 (2013).

31. Lin, M. T. et al. A procedure to introduce point mutations into the Rubisco large subunit gene in wild-type plants. The Plant Journal 106, 876–887 (2021).

32. Nakazato, I., Okuno, M., Itoh, T., Tsutsumi, N. & Arimura, S. Characterization and development of a plastid genome base editor, ptpTALECD. The Plant Journal 115, 1151–1162 (2023).

33. Clough, S. J. & Bent, A. F. Floral dip: a simplified method for Agrobacterium-mediated transformation of Arabidopsis thaliana. The plant journal 16, 735–743 (1998).

34. Shimada, T. L., Shimada, T. & Hara-Nishimura, I. A rapid and non-destructive screenable marker, FAST, for identifying transformed seeds of Arabidopsis thaliana. The Plant Journal 61, 519–528 (2010).

35. Nakazato, I. et al. Resistance to the herbicide metribuzin conferred to *Arabidopsis thaliana* by targeted base editing of the chloroplast genome. Plant Biotechnology Journal pbi.14490 (2024) doi:10.1111/pbi.14490.

36. Ishikawa, C., Hatanaka, T., Misoo, S., Miyake, C. & Fukayama, H. Functional incorporation of sorghum small subunit increases the catalytic turnover rate of Rubisco in transgenic rice. Plant Physiology 156, 1603–1611 (2011).

37. Yamori, W., Nagai, T. & Makino, A. The rate-limiting step for CO_2_ assimilation at different temperatures is influenced by the leaf nitrogen content in several C_3_ crop species. Plant Cell & Environment 34, 764–777 (2011).

38. Qu, Y. et al. Overexpression of both Rubisco and Rubisco activase rescues rice photosynthesis and biomass under heat stress. Plant Cell & Environment 44, 2308–2320 (2021).

39. Porra, R. J., Thompson, W. A. & Kriedemann, P. E. Determination of accurate extinction coefficients and simultaneous equations for assaying chlorophylls a and b extracted with four different solvents: verification of the concentration of chlorophyll standards by atomic absorption spectroscopy. Biochimica et Biophysica Acta (BBA)-Bioenergetics 975, 384–394 (1989).

40. Ishikawa, C., Hatanaka, T., Misoo, S. & Fukayama, H. Screening of high kcat Rubisco among Poaceae for improvement of photosynthetic CO2 assimilation in rice. Plant Production Science 12, 345–350 (2009).

41. Matsumura, H. et al. Crystal structure of rice Rubisco and implications for activation induced by positive effectors NADPH and 6-phosphogluconate. Journal of molecular biology 422, 75–86 (2012).

42. Mastronarde, D. N. Automated electron microscope tomography using robust prediction of specimen movements. Journal of structural biology 152, 36–51 (2005).

43. Yonekura, K., Maki-Yonekura, S., Naitow, H., Hamaguchi, T. & Takaba, K. Machine learning-based real-time object locator/evaluator for cryo-EM data collection. Communications Biology 4, 1044 (2021).

44. Zivanov, J. et al. New tools for automated high-resolution cryo-EM structure determination in RELION-3. elife 7, e42166 (2018).

45. Tegunov, D. & Cramer, P. Real-time cryo-electron microscopy data preprocessing with Warp. Nature methods 16, 1146–1152 (2019).

46. Pettersen, E. F. et al. UCSF Chimera—A visualization system for exploratory research and analysis. J Comput Chem 25, 1605–1612 (2004).

47. Emsley, P., Lohkamp, B., Scott, W. G. & Cowtan, K. Features and development of Coot. Acta Crystallographica Section D: Biological Crystallography 66, 486–501 (2010).

48. Adams, P. D. et al. PHENIX: a comprehensive Python-based system for macromolecular structure solution. Acta Crystallographica Section D: Biological Crystallography 66, 213–221 (2010).

49. Chen, V. B. et al. MolProbity: all-atom structure validation for macromolecular crystallography. Acta Crystallographica Section D: Biological Crystallography 66, 12–21 (2010).

50. Schindelin, J., Rueden, C. T., Hiner, M. C. & Eliceiri, K. W. The ImageJ ecosystem: An open platform for biomedical image analysis. Molecular Reproduction Devel 82, 518–529 (2015).

51. Larkin, M. A. et al. Clustal W and Clustal X version 2.0. bioinformatics 23, 2947–2948 (2007).

52. Robert, X. & Gouet, P. Deciphering key features in protein structures with the new ENDscript server. Nucleic acids research 42, W320–W324 (2014).

