## Extended Data for "Chloroplast genome editing of Rubisco boosts photosynthesis and plant growth"

**Extended Data Table 1: CryoEM data collection, refinement and validation statistics 1**

| Dataset | Col-0 Rubisco (Three classes merged with D4 symmetry) | M309I Rubisco (Three classes merged with D4 symmetry) | D397N Rubisco (Three classes merged with D4 symmetry) |
|---|---|---|---|
| EMDB accession code | EMD-62824 | EMD-62829 | EMD-62834 |
| PDB accession code | 9L58 | 9L5D | 9L5I |
| **Data collection and processing** | | | |
| Magnification | 60,000 | 60,000 | 60,000 |
| Voltage (kV) | 300 | 300 | 300 |
| Electron exposure ($e^{-}$/Å$^2$) | 80 | 80 | 80 |
| Defocus range (μm) | −0.5 to −2.0 | −0.5 to −2.0 | −0.5 to −2.0 |
| Pixel size (Å) | 0.878 | 0.878 | 0.878 |
| Symmetry imposed | D4 | D4 | D4 |
| Imported movies (no.) | 2,732 | 3,258 | 3,509 |
| Initial particle images (no.) | 1,487,227 | 2,074,623 | 2,302,408 |
| Final particle images (no.) | 737,173 | 1,170,331 | 904,485 |
| Map resolution (Å) | 1.76 | 1.75 | 1.74 |
| FSC threshold | 0.143 | 0.143 | 0.143 |
| Local resolution range (Å) | 1.74-2.48 | 1.74-2.21 | 1.74-2.39 |
| **Refinement** | | | |
| Initial model used (PDB code) | 6KYJ | Col-0 Rubisco (Three classes merged with D4 symmetry) | Col-0 Rubisco (Three classes merged with D4 symmetry) |
| Model resolution (Å) | 1.7/1.7/1.8 | 1.7/1.7/1.6 | 1.7/1.7/1.7 |
| FSC threshold | 0/0.143/0.5 | 0/0.143/0.5 | 0/0.143/0.5 |
| Model resolution (Å) | 1.76 | 1.75 | 1.74 |
| Map sharpening B factor (Å$^2$) | -44.23 | -45.51 | -42.04 |
| Model vs. Data CC (mask) | 0.87 | 0.77 | 0.87 |
| (volume) | 0.86 | 0.78 | 0.86 |
| Model composition | | | |
| Non-hydrogen atoms | 38,224 | 37,680 | 37,864 |
| Protein residues | 4,528 | 4,496 | 4,496 |
| Waters | 2,008 | 1,896 | 1,848 |
| Ligands | 16 ($SO_4$) | 8 ($SO_4$) | 8 ($SO_4$) |
| R.m.s. deviations | | | |
| Bond lengths (Å) | 0.007 | 0.006 | 0.004 |
| Bond angles (°) | 0.843 | 0.839 | 0.699 |
| Validation | | | |
| MolProbity score | 1.68 | 1.77 | 1.53 |
| Clashscore | 9.11 | 13.76 | 6.63 |
| Rotamer outliers (%) | 0.00 | 0.90 | 1.14 |
| Ramachandran plot | | | |
| Favored (%) | 96.84 | 97.35 | 97.37 |
| Allowed (%) | 3.16 | 2.65 | 2.63 |
| Outliers (%) | 0.00 | 0.00 | 0.00 |

**Extended Data Table 2: CryoEM data collection, refinement and validation statistics 2**

| Dataset | Col-0 Rubisco (Three classes merged with C1 symmetry) | M309I Rubisco (Three classes merged with C1 symmetry) | D397N Rubisco (Three classes merged with C1 symmetry) |
|---|---|---|---|
| EMDB accession code | EMD-62823 | EMD-62829 | EMD-62833 |
| PDB accession code | 9L57 | 9L5D | 9L5H |
| **Data collection and processing** | | | |
| Magnification | 60,000 | 60,000 | 60,000 |
| Voltage (kV) | 300 | 300 | 300 |
| Electron exposure ($e^{-}/Å^{2}$) | 80 | 80 | 80 |
| Defocus range (μm) | −0.5 to −2.0 | −0.5 to −2.0 | −0.5 to −2.0 |
| Pixel size (Å) | 0.878 | 0.878 | 0.878 |
| Symmetry imposed | C1 | C1 | C1 |
| Imported movies (no.) | 2,732 | 3,258 | 3,509 |
| Initial particle images (no.) | 1,487,227 | 2,074,623 | 2,302,408 |
| Final particle images (no.) | 737,666 | 1,170,331 | 904,485 |
| Map resolution (Å) | 1.95 | 1.85 | 1.87 |
| FSC threshold | 0.143 | 0.143 | 0.143 |
| Local resolution range (Å) | 1.87-2.98 | 1.77-2.67 | 1.79-4.24 |
| **Refinement** | | | |
| Initial model used (PDB code) | Col-0 Rubisco (Three classes merged with D4 symmetry) | M309I Rubisco (Three classes merged with D4 symmetry) | D397N Rubisco (Three classes merged with D4 symmetry) |
| Model resolution (Å) | 1.8/1.9/1.9 | 1.7/1.8/1.8 | 1.7/1.8/1.8 |
| FSC threshold | 0/0.143/0.5 | 0/0.143/0.5 | 0/0.143/0.5 |
| Model resolution (Å) | 1.95 | 1.85 | 1.87 |
| Map sharpening B factor ($Å^{2}$) | -41.76 | -39.61 | -37.52 |
| Model vs. Data CC (mask) | 0.88 | 0.87 | 0.89 |
| (volume) | 0.86 | 0.86 | 0.88 |
| Model composition | | | |
| Non-hydrogen atoms | 37,996 | 37,463 | 37,692 |
| Protein residues | 4,528 | 4,496 | 4,496 |
| Waters | 1,788 | 1,679 | 1,676 |
| Ligands | 16 ($SO_4$) | 8 ($SO_4$) | 8 ($SO_4$) |
| R.m.s. deviations | | | |
| Bond lengths (Å) | 0.005 | 0.006 | 0.007 |
| Bond angles (°) | 0.713 | 0.790 | 0.856 |
| Validation | | | |
| MolProbity score | 1.69 | 1.59 | 1.64 |
| Clashscore | 10.36 | 10.93 | 8.84 |
| Rotamer outliers (%) | 1.02 | 0.53 | 1.16 |
| Ramachandran plot | | | |
| Favored (%) | 97.20 | 97.86 | 97.44 |
| Allowed (%) | 2.80 | 2.14 | 2.56 |
| Outliers (%) | 0.00 | 0.00 | 0.00 |

**Extended Data Table 3: CryoEM data collection, refinement and validation statistics 3**

(Class1)

| Dataset | Col-0 Rubisco (Class1 with C1 symmetry) | M309I Rubisco (Class1 with C1 symmetry) | D397N Rubisco (Class1 with C1 symmetry) |
|---|---|---|---|
| EMDB accession code | EMD-62825 | EMD-62830 | EMD-62835 |
| PDB accession code | 9L59 | 9L5E | 9L5J |
| **Data collection and processing** | | | |
| Magnification | 60,000 | 60,000 | 60,000 |
| Voltage (kV) | 300 | 300 | 300 |
| Electron exposure ($e^{-}/Å^{2}$) | 80 | 80 | 80 |
| Defocus range (μm) | −0.5 to −2.0 | −0.5 to −2.0 | −0.5 to −2.0 |
| Pixel size (Å) | 0.878 | 0.878 | 0.878 |
| Symmetry imposed | C1 | C1 | C1 |
| Imported movies (no.) | 2,732 | 3,258 | 3,509 |
| Initial particle images (no.) | 1,487,227 | 2,074,623 | 2,302,408 |
| Final particle images (no.) | 89,314 | 338,656 | 258,006 |
| Map resolution (Å) | 2.42 | 2.01 | 2.10 |
| FSC threshold | 0.143 | 0.143 | 0.143 |
| Local resolution range (Å) | 2.22-4.26 | 1.90-3.14 | 2.01-3.42 |
| **Refinement** | | | |
| Initial model used (PDB code) | Col-0 Rubisco (Three classes merged with D4 symmetry) | M309I Rubisco (Three classes merged with D4 symmetry) | D397N Rubisco (Three classes merged with D4 symmetry) |
| Model resolution (Å) | 2.1/2.2/2.3 | 1.8/1.8/2.0 | 1.9/2.0/2.2 |
| FSC threshold | 0/0.143/0.5 | 0/0.143/0.5 | 0/0.143/0.5 |
| Model resolution (Å) | 2.42 | 2.01 | 2.10 |
| Map sharpening B factor ($Å^{2}$) | -39.14 | -38.79 | -49.86 |
| Model vs. Data CC (mask) | 0.88 | 0.87 | 0.88 |
| (volume) | 0.85 | 0.85 | 0.86 |
| Model composition | | | |
| Non-hydrogen atoms | 38,173 | 37,433 | 37,637 |
| Protein residues | 4,528 | 4,496 | 4,496 |
| Waters | 1,957 | 1,649 | 1,621 |
| Ligands | 16 ($SO_4$) | 8 ($SO_4$) | 8 ($SO_4$) |
| R.m.s. deviations | | | |
| Bond lengths (Å) | 0.003 | 0.005 | 0.006 |
| Bond angles (°) | 0.567 | 0.744 | 0.711 |
| Validation | | | |
| MolProbity score | 1.83 | 1.64 | 1.82 |
| Clashscore | 10.44 | 10.72 | 8.86 |
| Rotamer outliers (%) | 2.02 | 0.85 | 1.98 |
| Ramachandran plot | | | |
| Favored (%) | 97.71 | 97.55 | 97.39 |
| Allowed (%) | 2.20 | 2.43 | 2.61 |
| Outliers (%) | 0.09 | 0.02 | 0 |

**Extended Data Table 4: CryoEM data collection, refinement and validation statistics 4**

（Class2）

| Dataset | Col-0 Rubisco (Class2 with C1 symmetry) | M309I Rubisco (Class2 with C1 symmetry) | D397N Rubisco (Class2 with C1 symmetry) |
|---|---|---|---|
| EMDB accession code | EMD-62826 | EMD-62831 | EMD-62836 |
| PDB accession code | 9L5A | 9L5F | 9L5K |
| **Data collection and processing** | | | |
| Magnification | 60,000 | 60,000 | 60,000 |
| Voltage (kV) | 300 | 300 | 300 |
| Electron exposure ($e^-/Å^2$) | 80 | 80 | 80 |
| Defocus range (μm) | −0.5 to −2.0 | −0.5 to −2.0 | −0.5 to −2.0 |
| Pixel size (Å) | 0.878 | 0.878 | 0.878 |
| Symmetry imposed | C1 | C1 | C1 |
| Imported movies (no.) | 2,732 | 3,258 | 3,509 |
| Initial particle images (no.) | 1,487,227 | 2,074,623 | 2,302,408 |
| Final particle images (no.) | 274,725 | 578,985 | 306,772 |
| Map resolution (Å) | 2.02 | 1.95 | 2.01 |
| FSC threshold | 0.143 | 0.143 | 0.143 |
| Local resolution range (Å) | 1.89-3.24 | 1.86-2.91 | 1.89-3.31 |
| **Refinement** | | | |
| Initial model used (PDB code) | Col-0 Rubisco (Three classes merged with D4 symmetry) | M309I Rubisco (Three classes merged with D4 symmetry) | D397N Rubisco (Three classes merged with D4 symmetry) |
| Model resolution (Å) | 1.8/1.9/2.0 | 1.8/1.8/1.9 | 1.8/1.9/2.0 |
| FSC threshold | 0/0.143/0.5 | 0/0.143/0.5 | 0/0.143/0.5 |
| Model resolution (Å) | 2.02 | 1.95 | 2.01 |
| Map sharpening B factor ($Å^2$) | -38.77 | -39.90 | -35.24 |
| Model vs. Data CC (mask) | 0.87 | 0.87 | 0.88 |
| (volume) | 0.85 | 0.85 | 0.87 |
| Model composition | | | |
| Non-hydrogen atoms | 38,002 | 37,436 | 37,864 |
| Protein residues | 4,528 | 4,496 | 4,496 |
| Waters | 1,802 | 1,652 | 1,848 |
| Ligands | 16 ($SO_4$) | 8 ($SO_4$) | 8 ($SO_4$) |
| R.m.s. deviations | | | |
| Bond lengths (Å) | 0.003 | 0.003 | 0.008 |
| Bond angles (°) | 0.617 | 0.645 | 0.837 |
| Validation | | | |
| MolProbity score | 1.58 | 1.62 | 1.75 |
| Clashscore | 10.07 | 10.41 | 8.39 |
| Rotamer outliers (%) | 0.63 | 1.01 | 1.51 |
| Ramachandran plot | | | |
| Favored (%) | 97.75 | 97.62 | 97.10 |
| Allowed (%) | 2.22 | 2.32 | 2.90 |
| Outliers (%) | 0.02 | 0.07 | 0 |

**Extended Data Table 5: CryoEM data collection, refinement and validation statistics 5**
（Class3）

| Dataset | Col-0 Rubisco (Class3 with C1 symmetry) | M309I Rubisco (Class3 with C1 symmetry) | D397N Rubisco (Class3 with C1 symmetry) |
|---|---|---|---|
| EMDB accession code | EMD-62827 | EMD-62832 | EMD-62837 |
| PDB accession code | 9L5B | 9L5G | 9L5L |
| **Data collection and processing** | | | |
| Magnification | 60,000 | 60,000 | 60,000 |
| Voltage (kV) | 300 | 300 | 300 |
| Electron exposure ($e^{-}/Å^2$) | 80 | 80 | 80 |
| Defocus range (μm) | −0.5 to −2.0 | −0.5 to −2.0 | −0.5 to −2.0 |
| Pixel size (Å) | 0.878 | 0.878 | 0.878 |
| Symmetry imposed | C1 | C1 | C1 |
| Imported movies (no.) | 2,732 | 3,258 | 3,509 |
| Initial particle images (no.) | 1,487,227 | 2,074,623 | 2,302,408 |
| Final particle images (no.) | 353,069 | 252,690 | 339,707 |
| Map resolution (Å) | 2.08 | 2.04 | 2.01 |
| FSC threshold | 0.143 | 0.143 | 0.143 |
| Local resolution range (Å) | 1.98-3.28 | 1.93-3.15 | 1.89-3.33 |
| **Refinement** | | | |
| Initial model used (PDB code) | Col-0 Rubisco (Three classes merged with D4 symmetry) | M309I Rubisco (Three classes merged with D4 symmetry) | D397N Rubisco (Three classes merged with D4 symmetry) |
| Model resolution (Å) | 1.9/2.0/2.0 | 1.9/1.9/2.0 | 1.8/1.9/2.0 |
| FSC threshold | 0/0.143/0.5 | 0/0.143/0.5 | 0/0.143/0.5 |
| Model resolution (Å) | 2.08 | 2.04 | 2.01 |
| Map sharpening B factor ($Å^2$) | -43.05 | -39.56 | -35.62 |
| Model vs. Data CC (mask) | 0.89 | 0.86 | 0.80 |
| (volume) | 0.87 | 0.84 | 0.79 |
| Model composition | | | |
| Non-hydrogen atoms | 37,926 | 37,302 | 37,578 |
| Protein residues | 4,528 | 4,496 | 4,496 |
| Waters | 1,722 | 1,564 | 1,582 |
| Ligands | 16 ($SO_4$) | 8 ($SO_4$) | 8 ($SO_4$) |
| R.m.s. deviations | | | |
| Bond lengths (Å) | 0.004 | 0.006 | 0.004 |
| Bond angles (°) | 0.699 | 0.796 | 0.714 |
| Validation | | | |
| MolProbity score | 1.64 | 1.92 | 1.74 |
| Clashscore | 11.77 | 13.94 | 13.33 |
| Rotamer outliers (%) | 0.87 | 1.36 | 1.01 |
| Ramachandran plot | | | |
| Favored (%) | 97.73 | 97.05 | 97.44 |
| Allowed (%) | 2.25 | 2.95 | 2.54 |
| Outliers (%) | 0.02 | 0.00 | 0.02 |

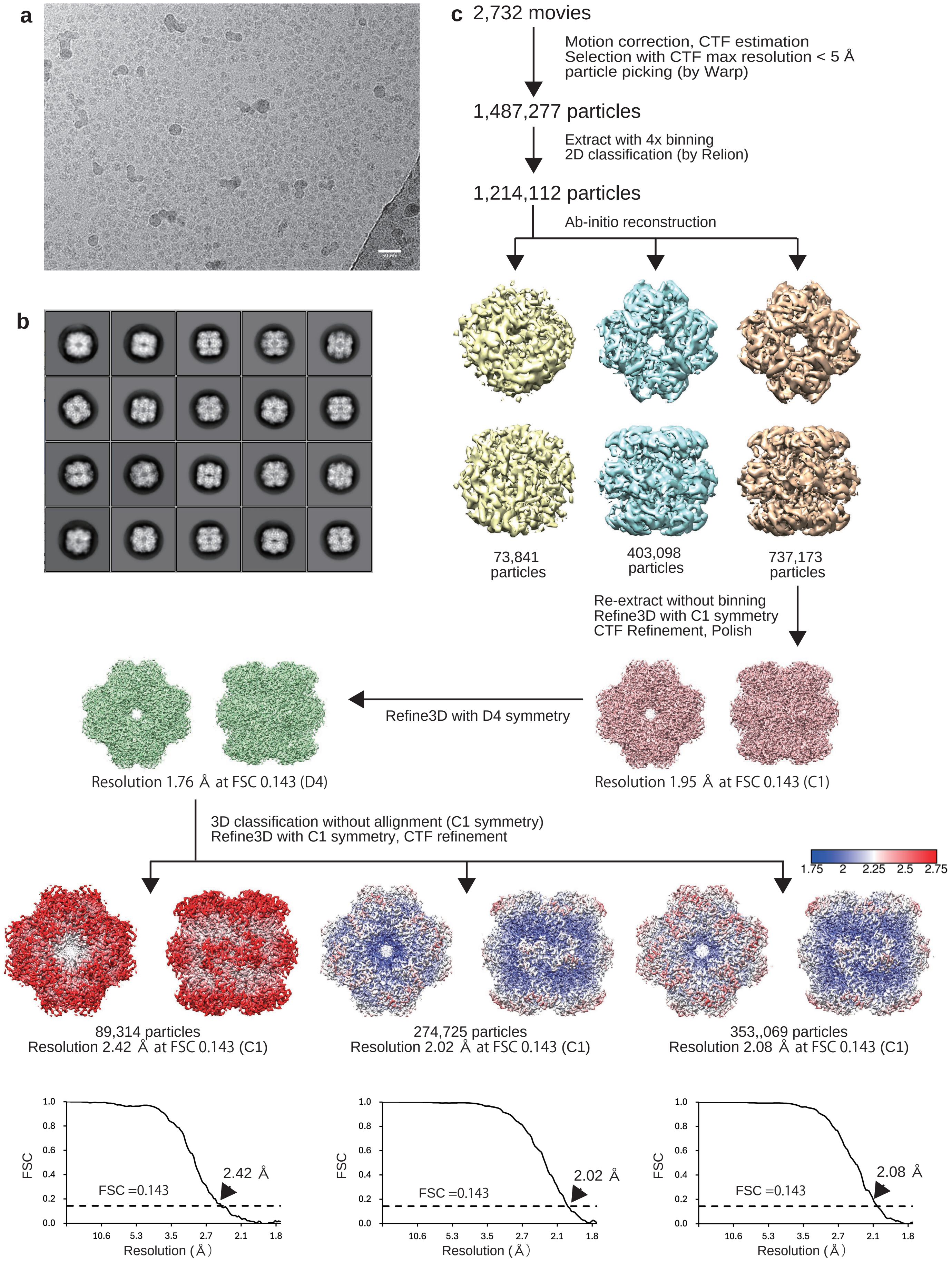

**Extended Data Fig. 1 Cryo-EM single-particle 3D image analysis of the Col-0 Rubisco. a**, Representative cryo-EM image of Col-0 Rubisco. Bar = 50 nm. **b**, Reference-free two-dimensional averages of Col-0 Rubisco. **c**, Work process of Cryo-EM single-particle 3D reconstruction of Col-0 Rubisco and Fourier Shell Correlation (FSC) curves of the final density maps of Col-0 Rubisco reconstructed with C1 symmetry. The local resolution of the maps is indicated by a color gradient ranging from blue (1.75 Å) to red (2.75 Å).

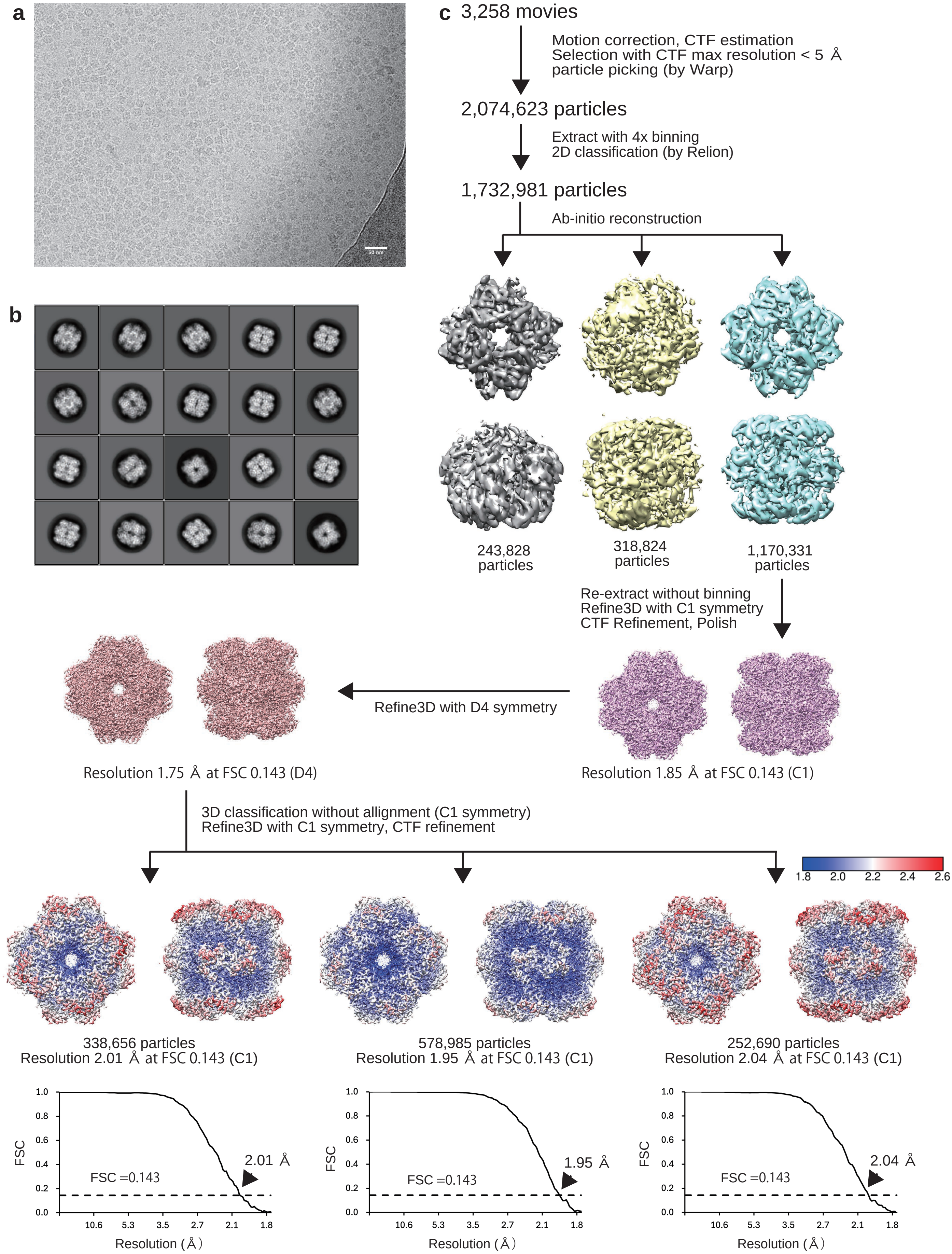

**Extended Data Fig. 2 Cryo-EM single-particle 3D image analysis of the M309I Rubisco. a**, Representative cryo-EM image of the M309I Rubisco. Bar = 50 nm. **b**, Reference-free two-dimensional averages of the M309I Rubisco. **c**, Work process of cryo-EM single-particle 3D reconstruction of the M309I Rubisco and Fourier Shell Correlation (FSC) curves of the final density maps of the M309I Rubisco reconstructed with C1 symmetry. The local resolution of the maps is indicated by a color gradient ranging from blue (1.8 Å) to red (2.6 Å).

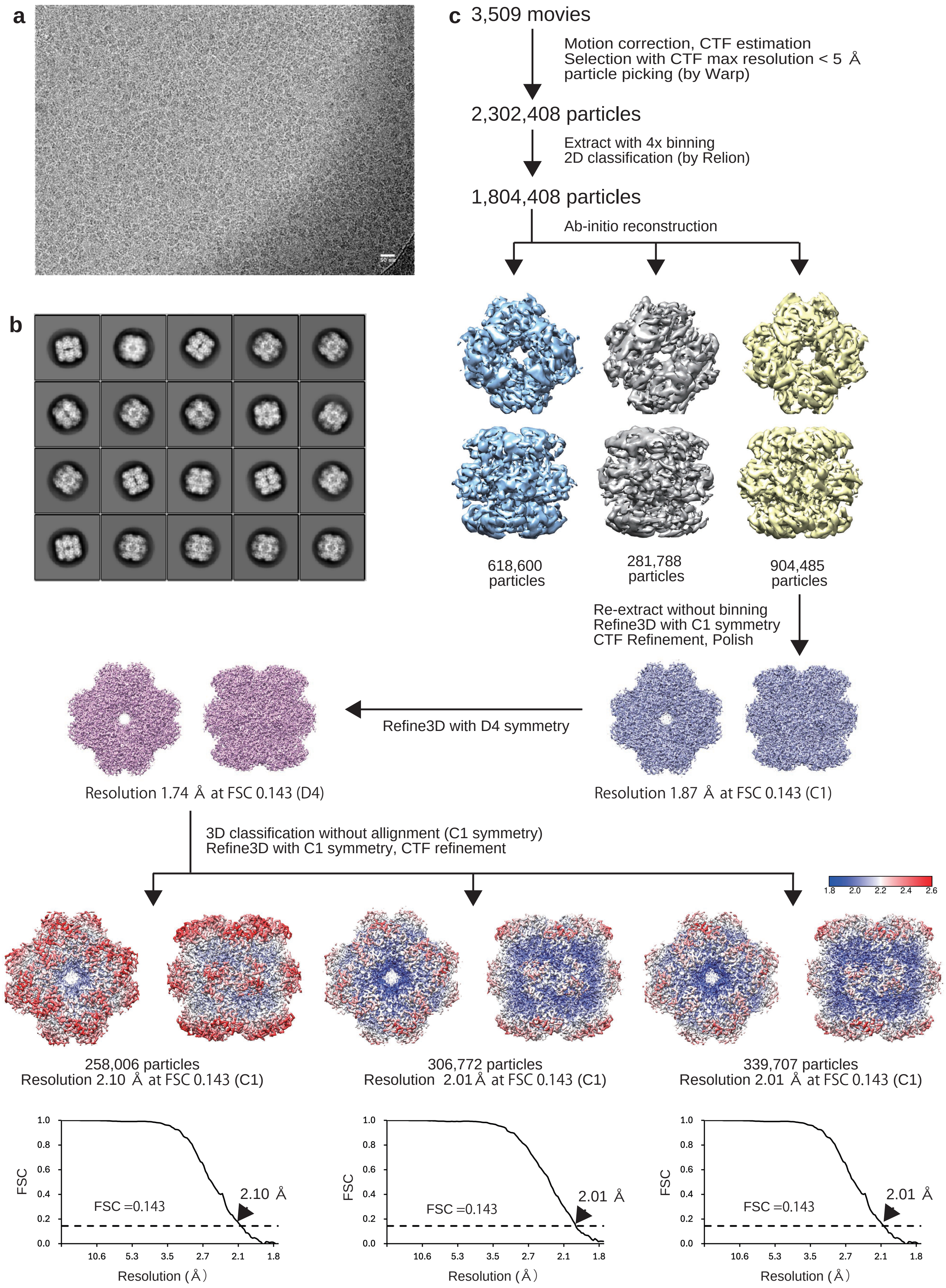

**Extended Data Fig. 3 Cryo-EM single-particle 3D image analysis of the D397N Rubisco. a**, Representative cryo-EM image of the D397N Rubisco. Bar = 50 nm. **b**, Reference-free two-dimensional averages of the D397N Rubisco. **c**, Work process of cryo-EM single-particle 3D reconstruction of the D397N Rubisco and Fourier Shell Correlation (FSC) curves of the final density maps of the D397N Rubisco reconstructed with C1 symmetry. The local resolution of the maps is indicated by a color gradient ranging from blue (1.8 Å) to red (2.6 Å).

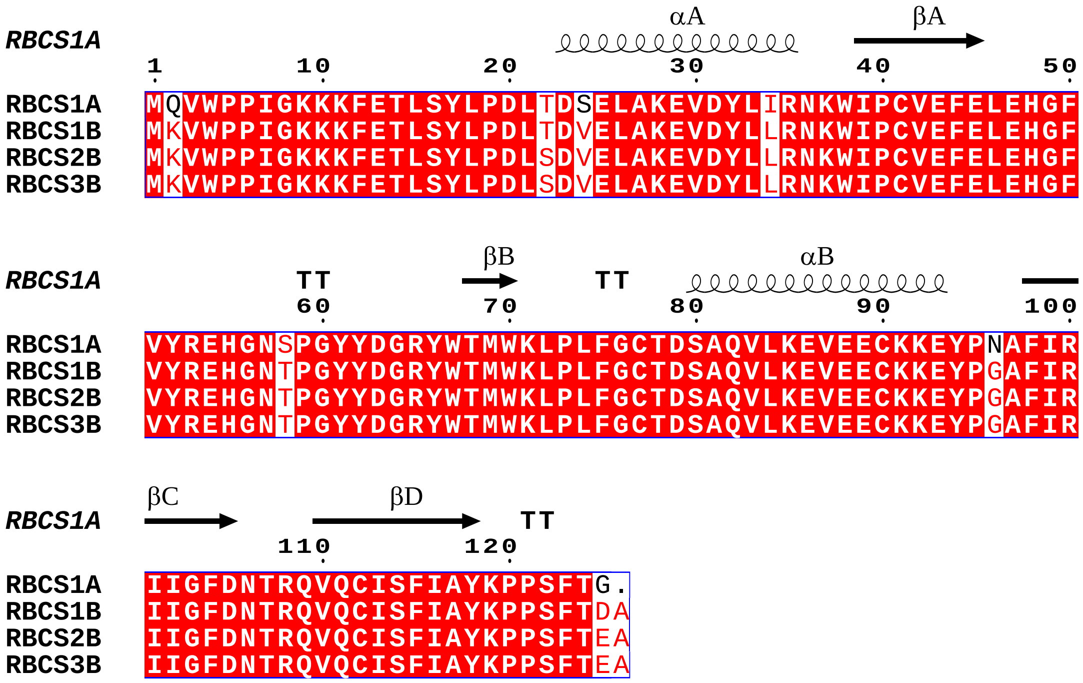

**Extended Data Fig. 4 Sequence alignment of RbcS isoforms in *A. thaliana***. The amino acid sequences of RBCS1A RBCS1B, RBCS2B, and RBCS3B are provided in GenBank accession number AEE34594.1, BAB09355.1, AAO29974.1, and AAL47390.1, respectively. Secondary structural elements are indicated above the amino acid sequence by arrows and coils for β-strands and α-helices, respectively. In addition, amino acids conserved across all four isoforms are highlighted with a red background.

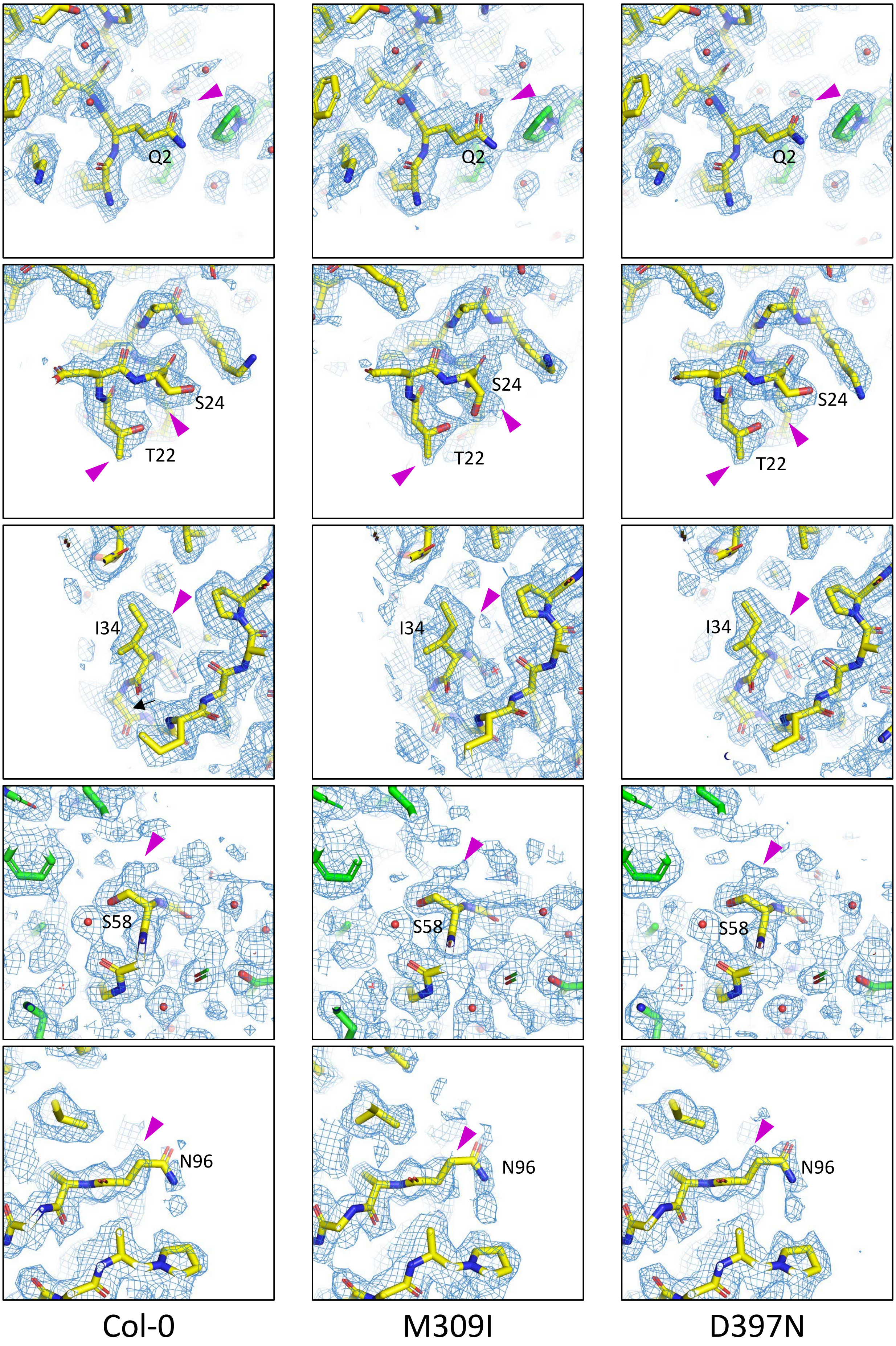

**Extended Data Fig. 5 Cryo-EM density maps of *A. thaliana* RbcS.** Cryo-EM density maps contoured at 1.5 σ around the amino acid residues that differ among RbcS isoforms. Maps of Col-0, M309I, and D397N are shown in the left, middle, and right panels, respectively. The amino acid residues are annotated based on the RBCS1A sequence (Extended Data Fig. 4). The magenta arrowheads show a mixture of RbcS isoforms.

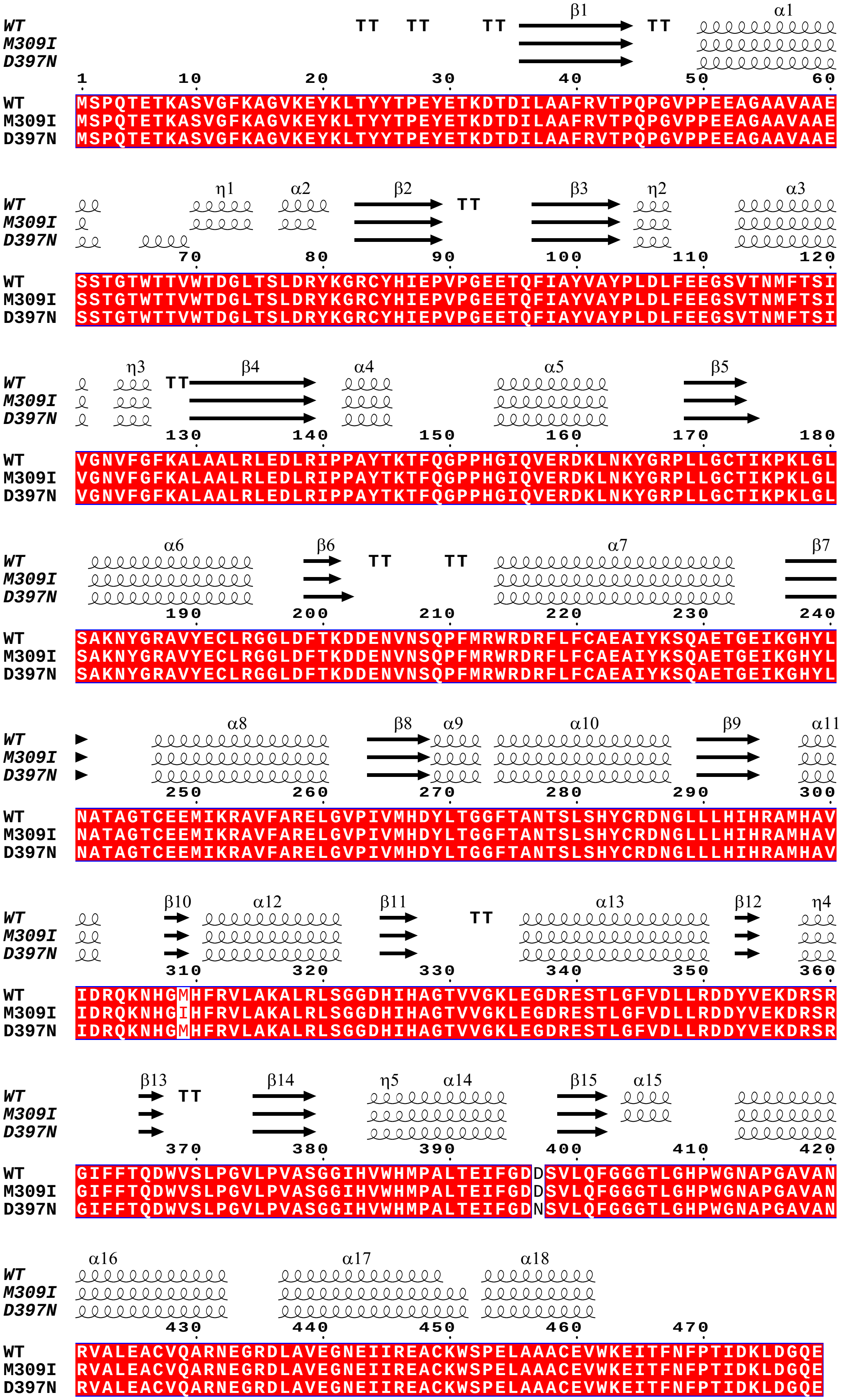

**Extended Data Fig. 6 The secondary structures of Col-0, M309I, and D397N RbcL of Rubisco.** The amino acid sequence of Col-0 is provided in GenBank accession number AAB68400.1. Secondary structural elements are indicated above the amino acid sequence by arrows and coils for β-strands and α-helices, respectively.
