## Supplementary Figures for "Chloroplast genome editing of Rubisco boosts photosynthesis and plant growth"

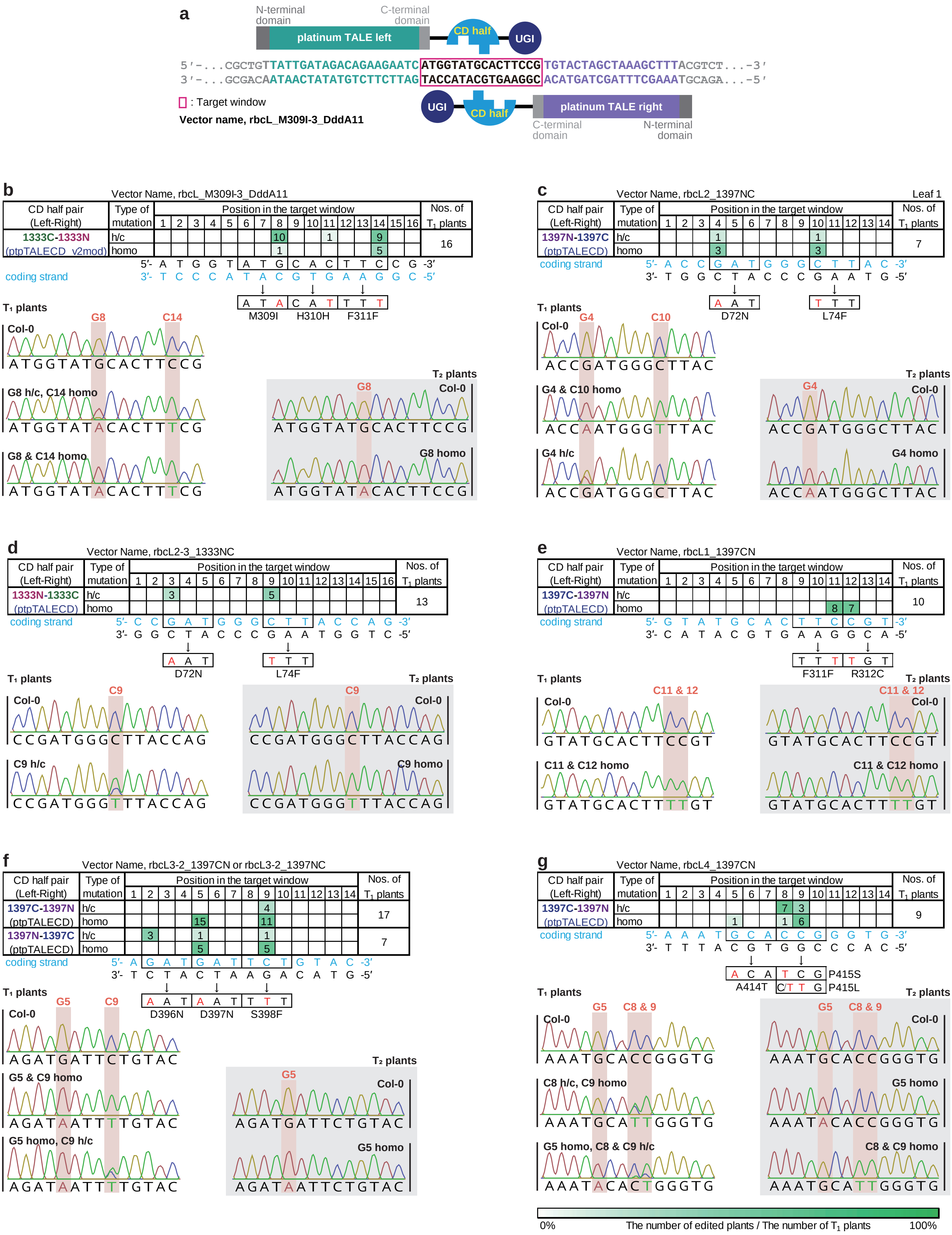


**Fig. S1 Base editing in the T$_1$ generation. a**, A schematic image of interaction between the base editor and their target sequence. platinum TALE, the DNA binding domain of platinum TALEN (Sakuma et al., 2013); CD, cytidine deaminase; UGI, uracil glycosylase inhibitor. **b-g**, Base editing efficiency in the T$_1$ generation and Sanger traces of T$_1$ and T$_2$ plants. The tables in the upper panel show which bases in the target window were edited, how frequently bases were edited, and how many plants underwent base editing. Genotypes were determined by Sanger sequencing, and the representative Sanger traces of T$_1$ and T$_2$ plants are shown in the lower left and right panels, respectively. Allele patterns of T$_2$ plants are shown in Table S2. The constructs were rbcL_M309I-3_DddA11 (b), rbcL2_1397NC (c), rbcL2-3_1333NC (d), rbcL1_1397CN (e), rbcL3-2_1397CN (or NC) (f), and rbcL4_1397CN (g). homo, homoplasmically edited; h/c, heteroplasmically or chimerically (not homoplasmically) edited.

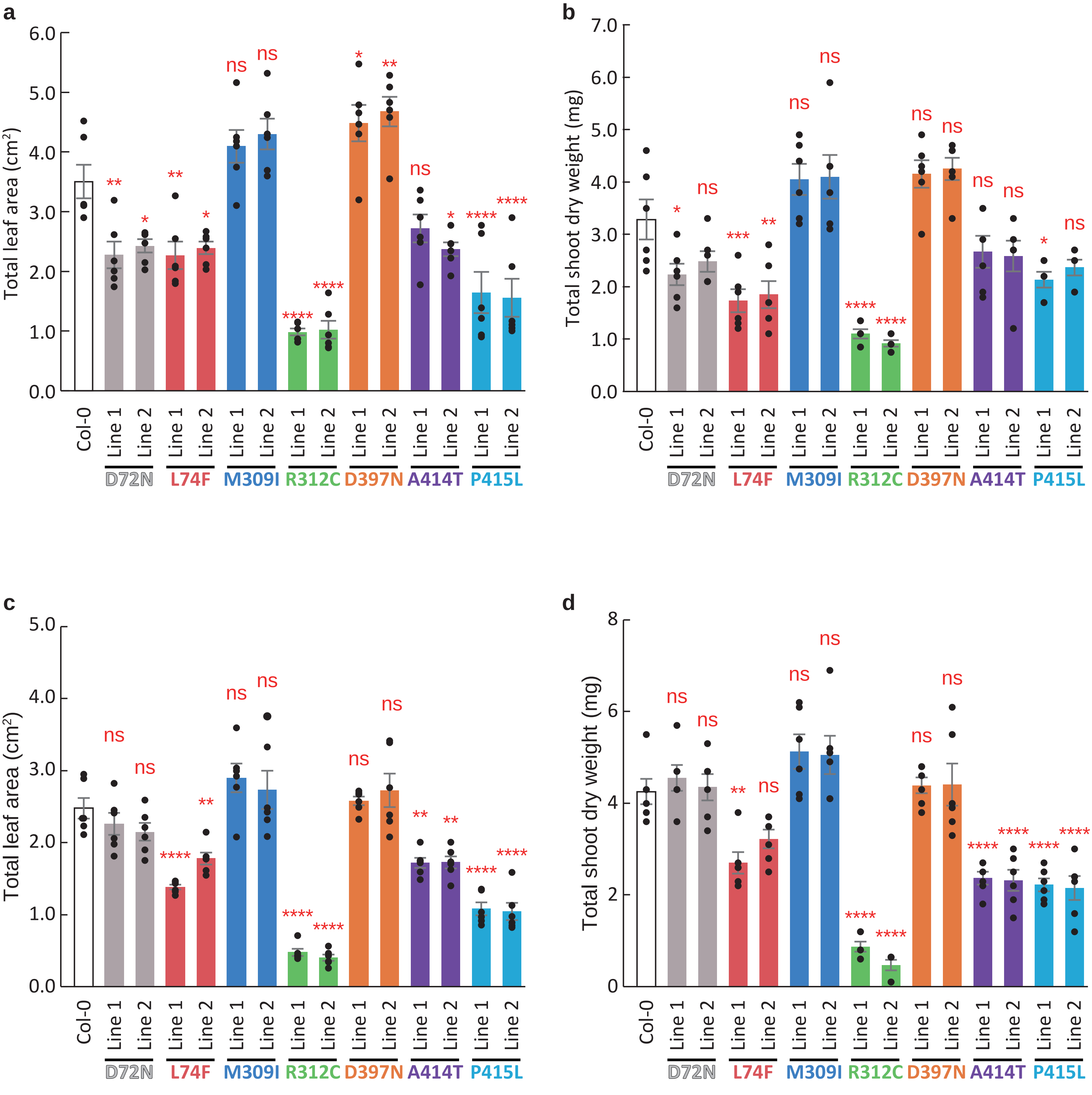


**Fig. S2 Plant growth at three weeks age in ambient (381 ppm±14) and high (549 ppm±23) $CO_2$ concentration.** **a, b**, Total leaf area (a), and total shoot dry weight (b) of plants cultivated under ambient $CO_2$ concentration at 24 days after sowing. **c, d**, Total leaf area (c), and total shoot dry weight (d) of plants cultivated under high $CO_2$ concentration at 25 days after sowing. Data is mean ± standard error. Individual data points were displayed as black dots. ns $p > 0.05$, * $p < 0.05$, ** $p < 0.01$, *** $p < 0.001$, **** $p < 0.0001$, n = 6 plants. Dunnett's two-tailed test for multiple comparisons.

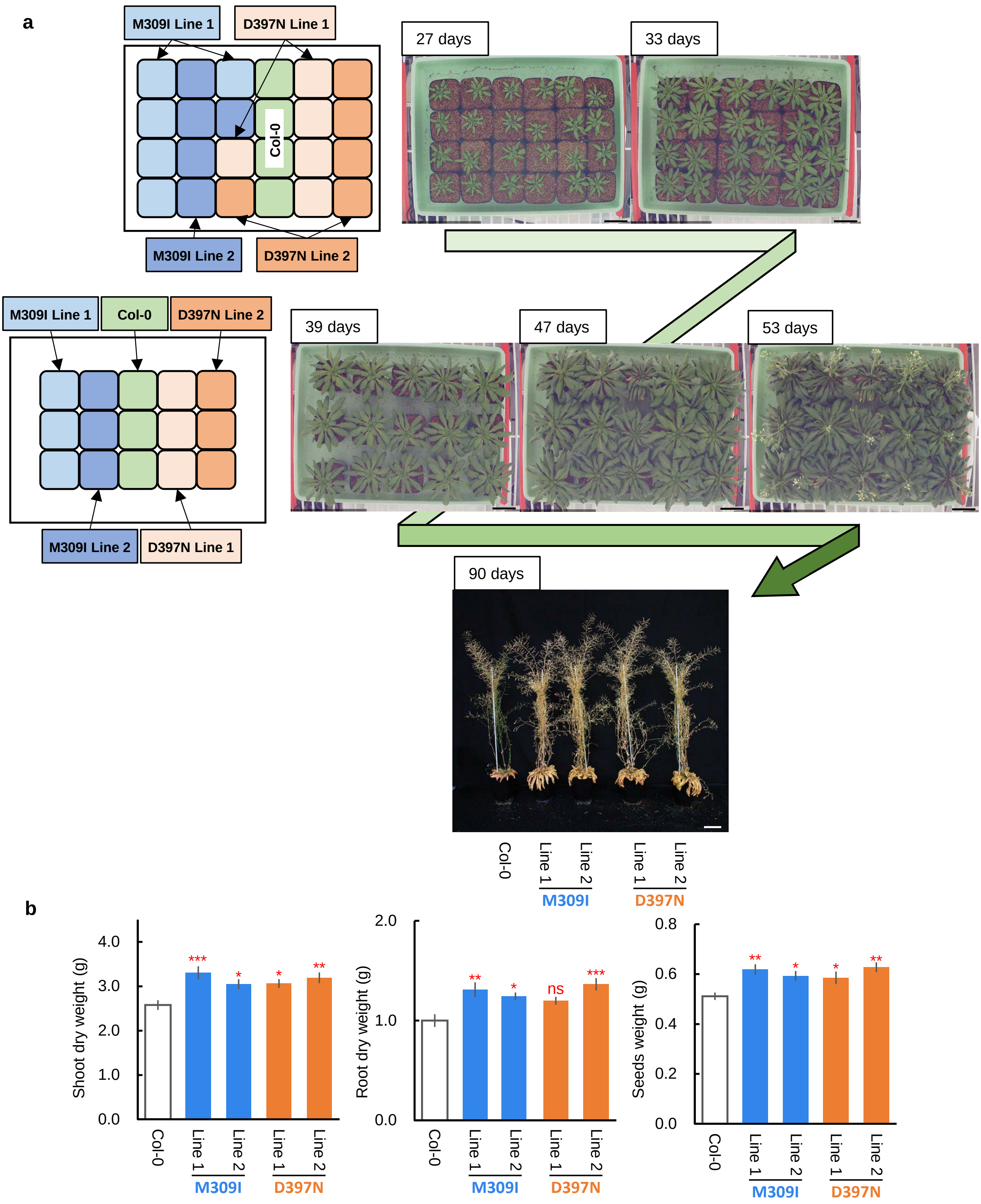


**Fig. S3 Plant growth in ambient $CO_2$ concentration (381 ppm±14). a**, Plant growth over a 90-day period, from seedling to harvest. No significant differences in bolting time were observed across all plants. Plants were grown with a daylight/dark period of 10/14 h, and photosynthetic photon flux density (PPFD) in the daylight period was set to 150 μmol photons $m^{-2}$ $s^{-1}$. Air temperature and relative humidity were maintained at 22°C and 60%, respectively. Bars = 5 cm. **b**, Weights of dried shoots and roots and seeds harvested at day 90. Data are means ± SE, n = 5 plants. ns $p$ > 0.05, * $p$ < 0.05, ** $p$ < 0.01, *** $p$ < 0.001, indicating significant differences of each column compared with Col-0. Dunnett's two-tailed test for multiple comparisons.

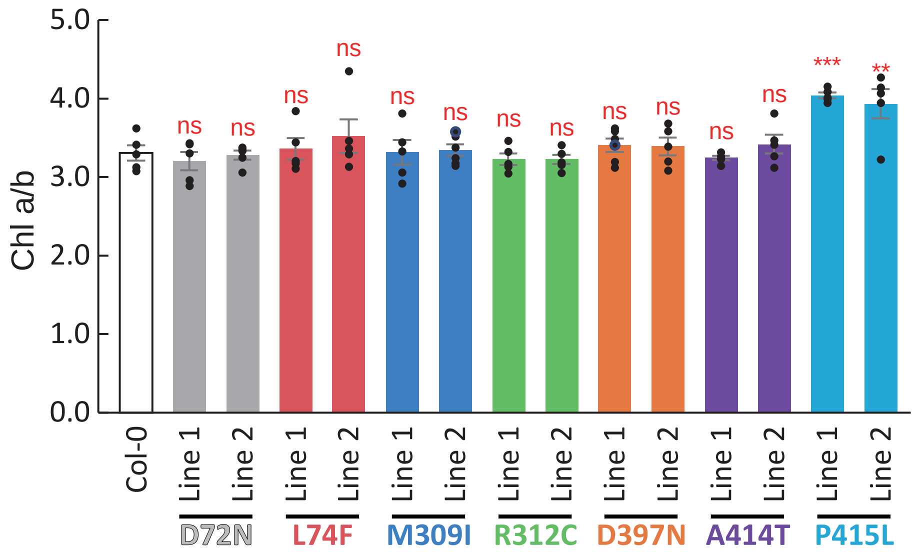


**Fig. S4 Chlorophyll a/b ratio in different lines.** Data is mean ± SE. ns $p > 0.05$, * $p < 0.05$, ** $p < 0.01$, *** $p < 0.001$, **** $p < 0.0001$, n = 6 plants. Dunnett's two-tailed test for multiple comparisons.

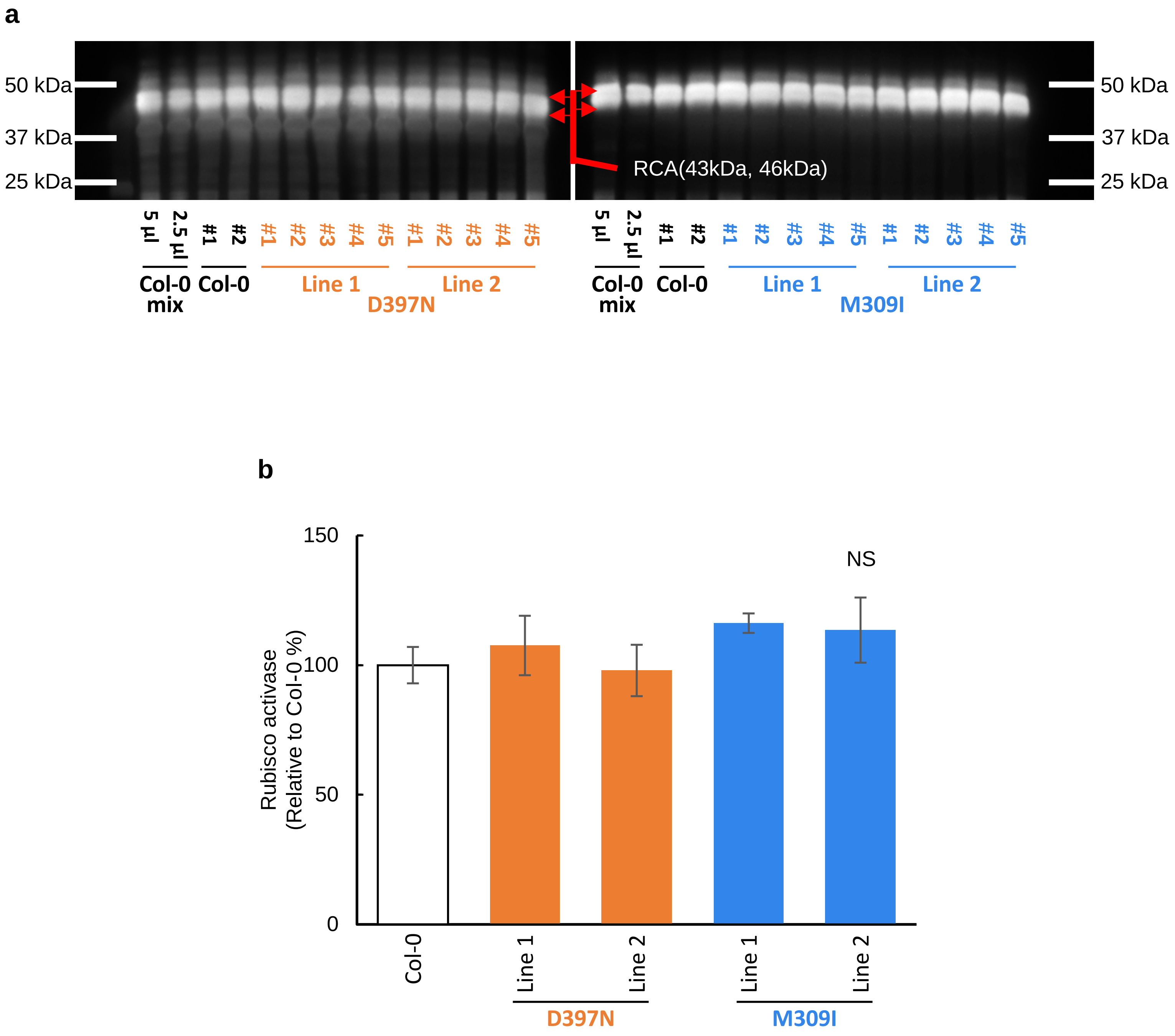


**Fig. S5 Analysis of Rubisco activase expression in M309I and D397N single mutants. a**, Western blots targeting Rubisco activase (RCA). Western blots were probed with polyclonal antibodies raised against rice Rubisco activase, and visualized using alkaline phosphatase conjugated to a secondary antibody. Two RCA isoforms, with molecular weights of 43 kDa and 46 kDa, were detected. Except for the lane labeled "Col-0 mix 2.5 µL", 5 µL of protein solution was loaded into each well. **b**, Relative amount of Rubisco activase against Col-0 (normalized to 100%) in different lines. Data are means ± SE, n = 4 (for Col-0) or 5 (for each different line) biological replicates. NS, not significant by Dunnett's two-tailed test for multiple comparisons.

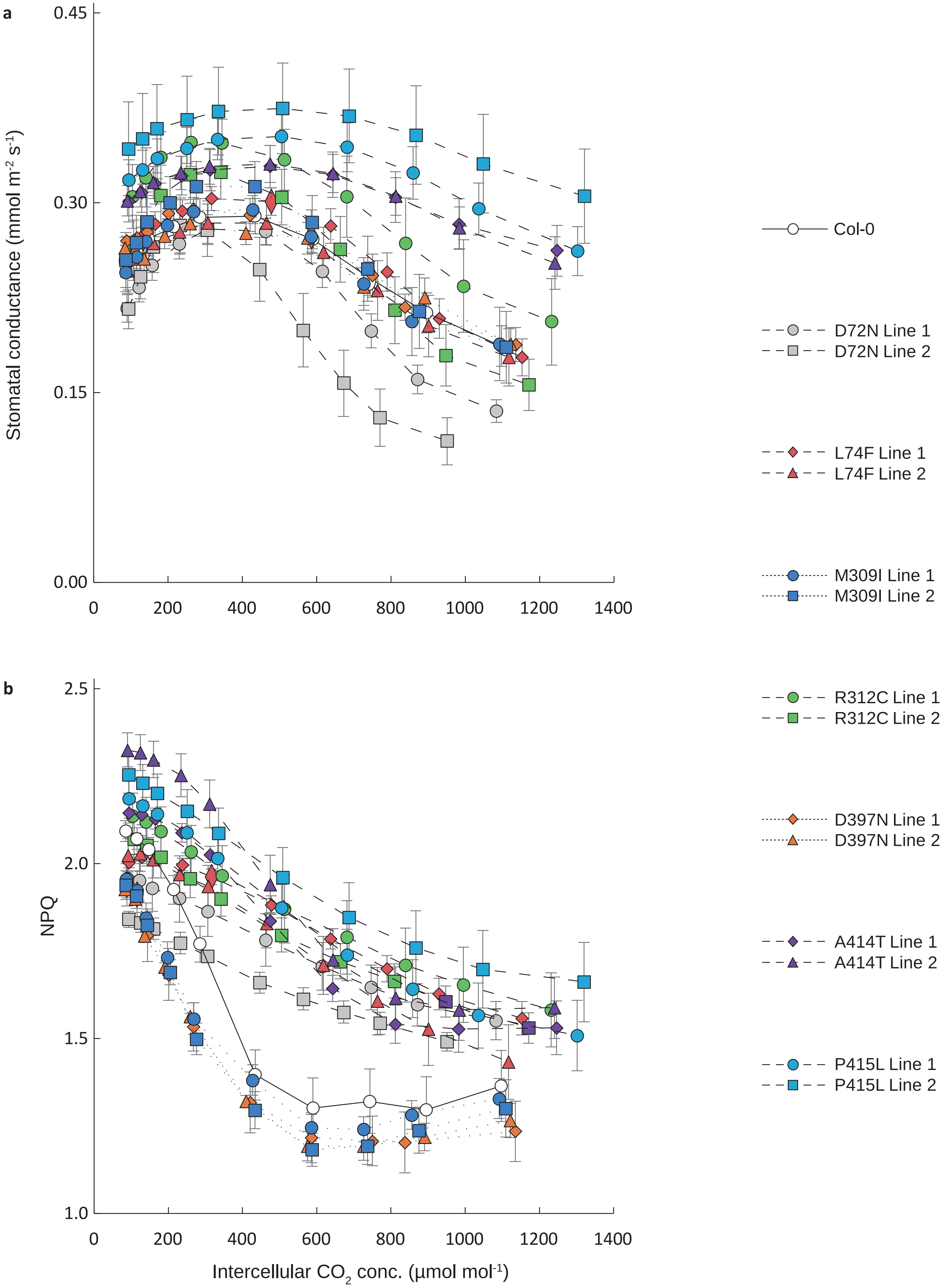


**Fig. S6 Photosynthesis parameters in various intercellular $CO_2$ concentrations of different lines. a**, Stomatal conductance of different lines in various intercellular $CO_2$ concentrations. **b**, NPQ of different lines in various intercellular $CO_2$ concentrations. Data is mean ± SE. Individual data points were displayed as black dots. ns $p > 0.05$, * $p < 0.05$, ** $p < 0.01$, *** $p < 0.001$, **** $p < 0.0001$, n = 6 plants. Dunnett's two-tailed test for multiple comparisons.

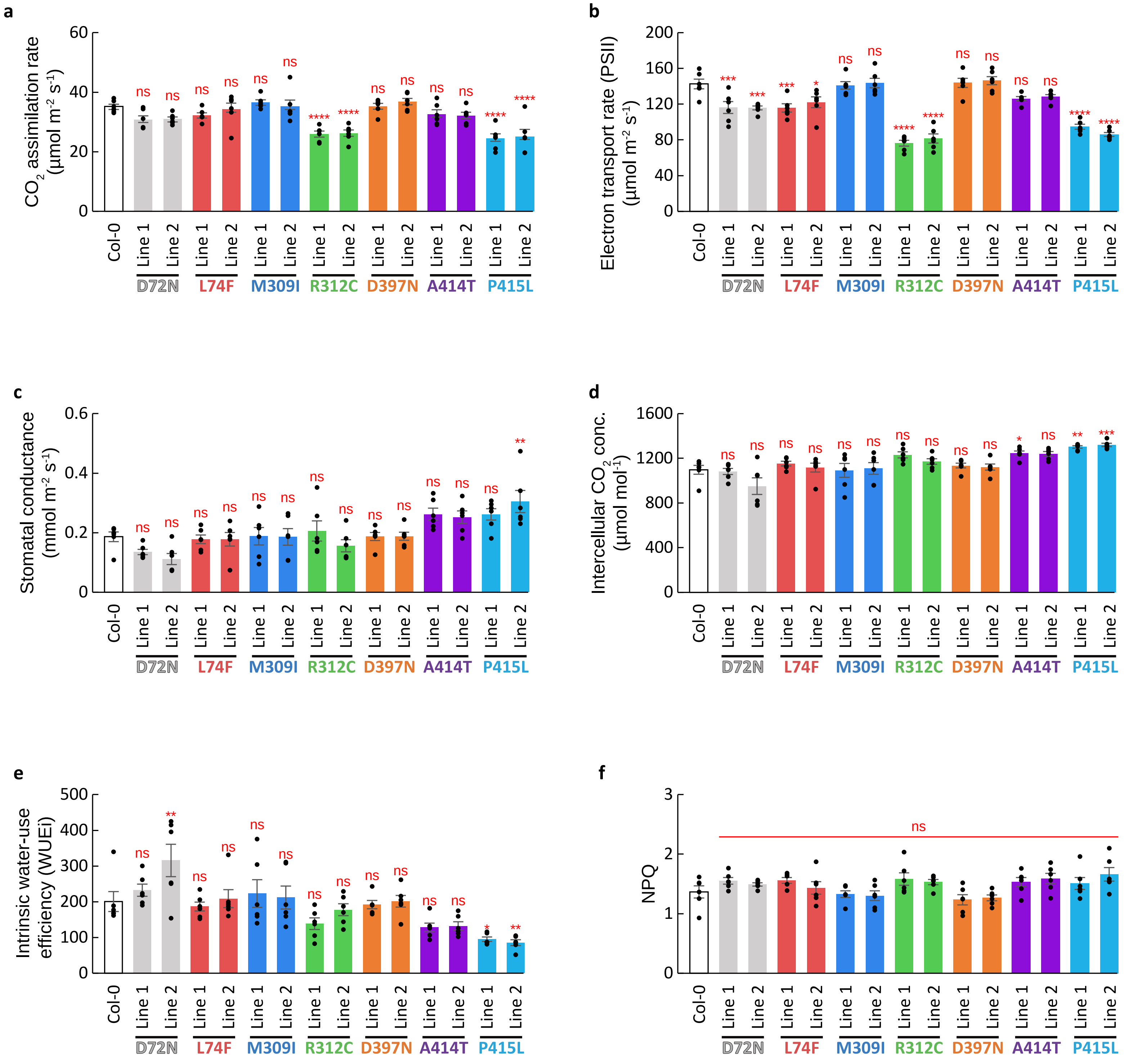


**Fig. S7 Photosynthesis parameters in 1200 ppm $CO_2$ of different lines.** **a**, $CO_2$ assimilation rate. **b**, Electron transport rate of PSII. **c**, Stomatal conductance. **d**, Intercellular $CO_2$ concentration. **e**, Intrinsic water-use efficiency (WUEi). **f**, NPQ. Data is mean ± SE. Individual data points were displayed as black dots. ns $p > 0.05$, * $p < 0.05$, ** $p < 0.01$, *** $p < 0.001$, **** $p < 0.0001$, n = 6 plants. Dunnett's two-tailed test for multiple comparisons.

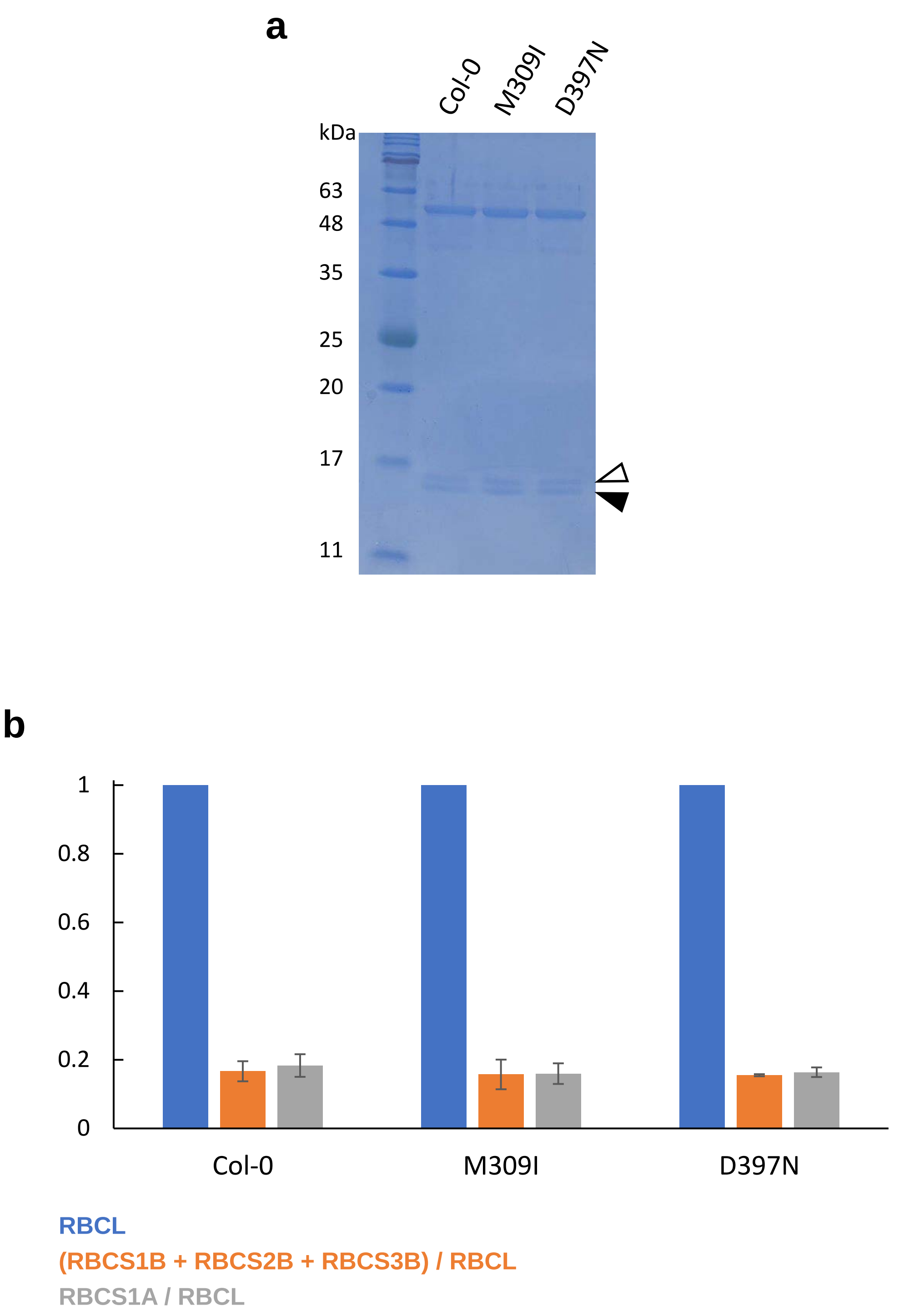


**Fig. S8 SDS-PAGE analysis of *A. thaliana* Rubisco purified from Col-0, M309I, and D397N leaves. a**, A representative SDS-PAGE image of Col-0, M309I, and D397N Rubisco. The white arrowhead indicates the RBCS1B, RBCS2B, and RBCS3B proteins, while the black arrowhead indicates RBCS1A, as reported by Izumi *et al* 2012 (Ref. 23). **b**, Ratios of RBCS protein band intensities to RBCL proteins. The blue bars represent the relative amount of RBCL (normalized to 1.0). The ratios of RBCS1B, RBCS2B, and RBCS3B proteins to RBCL are shown in orange, while the ratios of RBCS1A proteins to RBCL are shown in grey. Band intensities were quantified using ImageJ version 1.53q. Data represent the average of three independent experiments, with error bars indicating standard deviation.

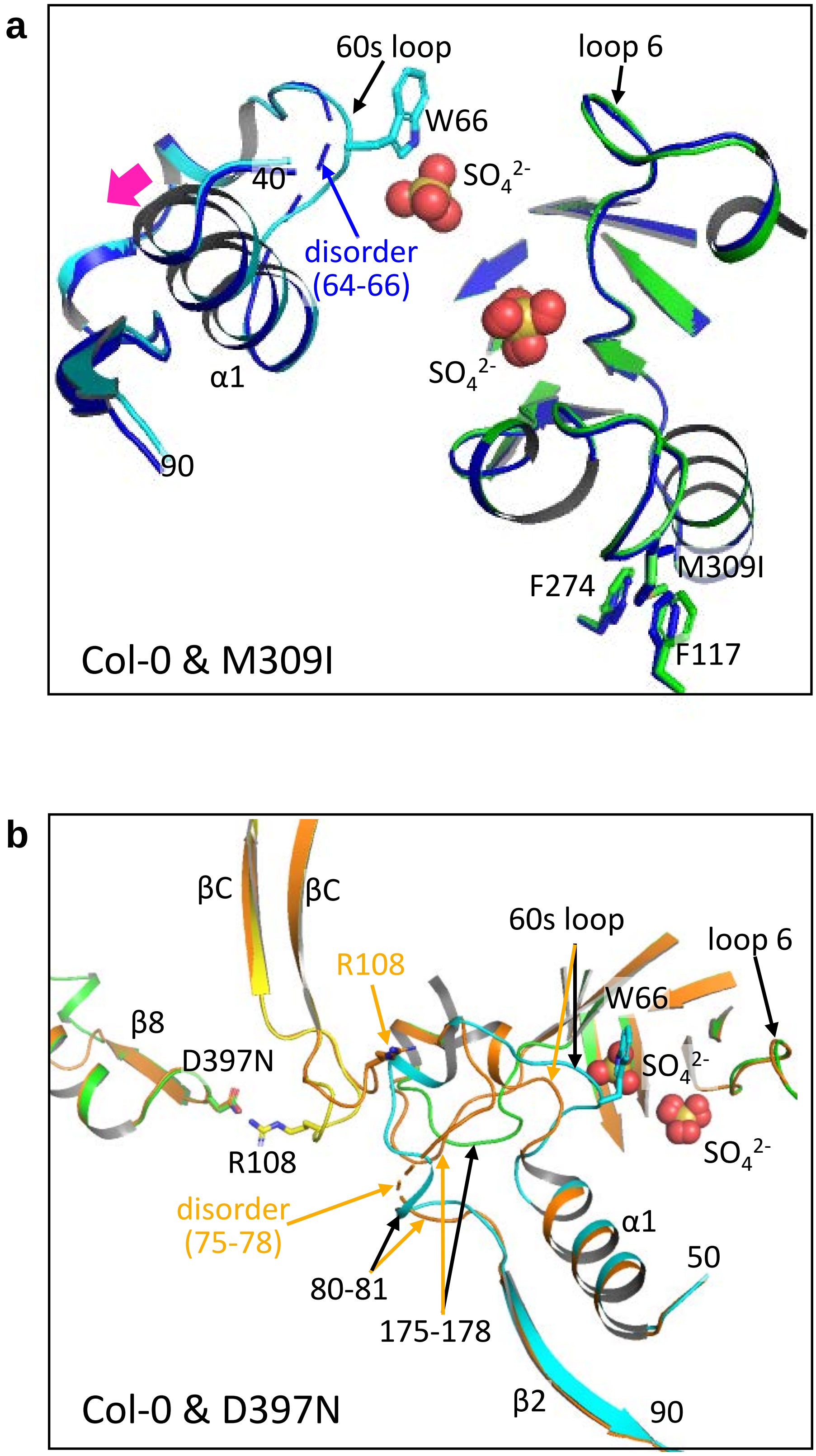


**Fig. S9 Superimposition of the active site structures of the *A. thaliana* Rubisco in complex with sulfate ions. a**, Rubisco structure around the 60s loop and loop 6 in Col-0 (green and cyan) and M309I (blue). The magenta arrow indicates a shift of approximately 0.5 Å in the α1 helix. Key interacting residues and regions are labeled. **b**, Rubisco structure around the 60s loop and loop 6 in Col-0 (green, cyan, and yellow) and D397N (orange). Black and orange arrows indicate the partial structures of Col-0 and D397N Rubisco, respectively.

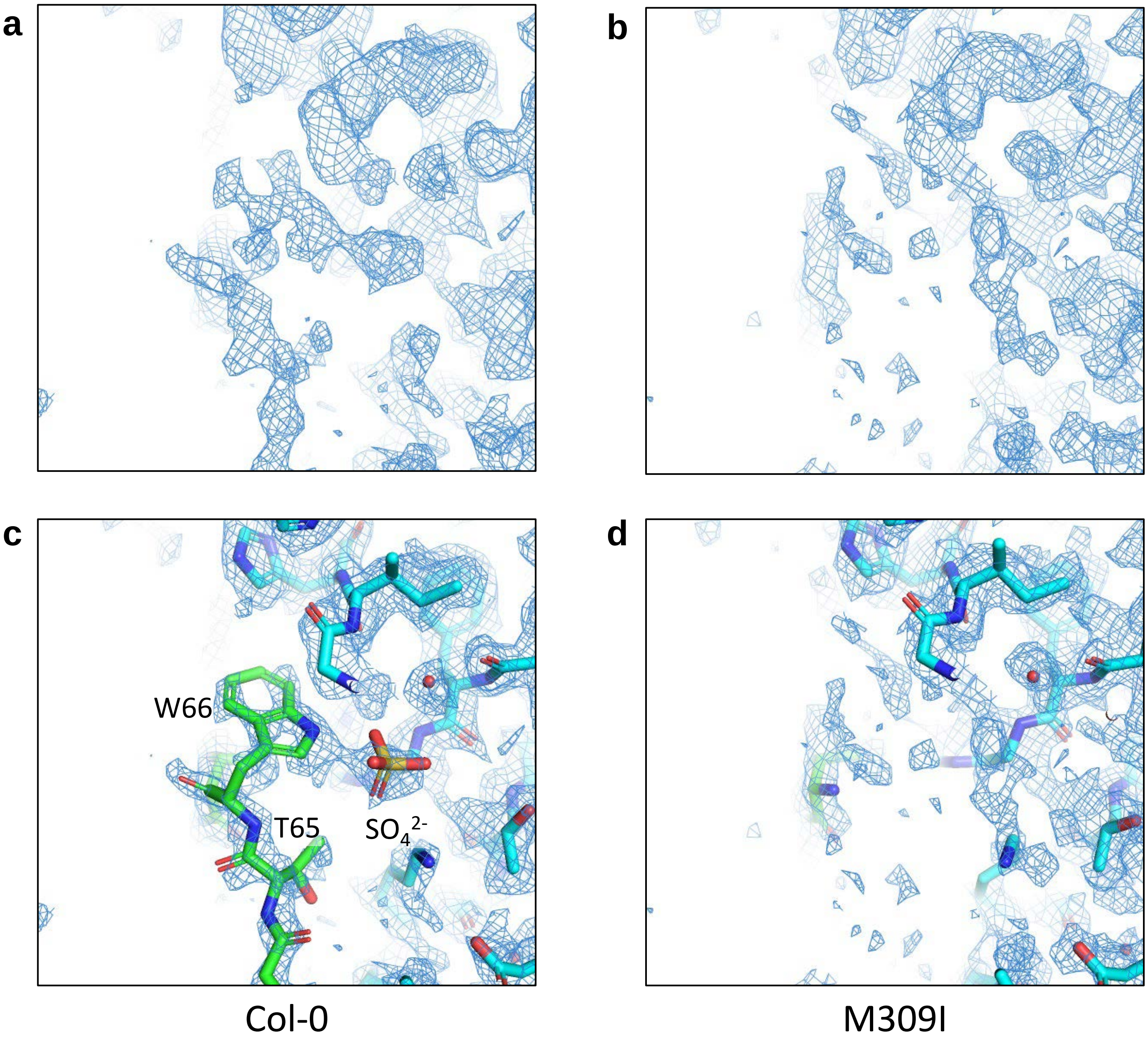


**Fig. S10 Differences in cryo-EM density maps around the 60s loop between Col-0 and M309I Rubisco. a, b**, The cryo-EM density maps contoured at 2.0 σ for Col-0 (a) and M309I (b). **c, d**, Fitness of residues in the 60s loop in Col-0 (c) and M309I (d). The cryo-EM density maps of Col-0 allowed fitting all portions of the 60s loop, while the region corresponding to residues 64–66 in this loop could not be fitted due to lower cryo-EM density.

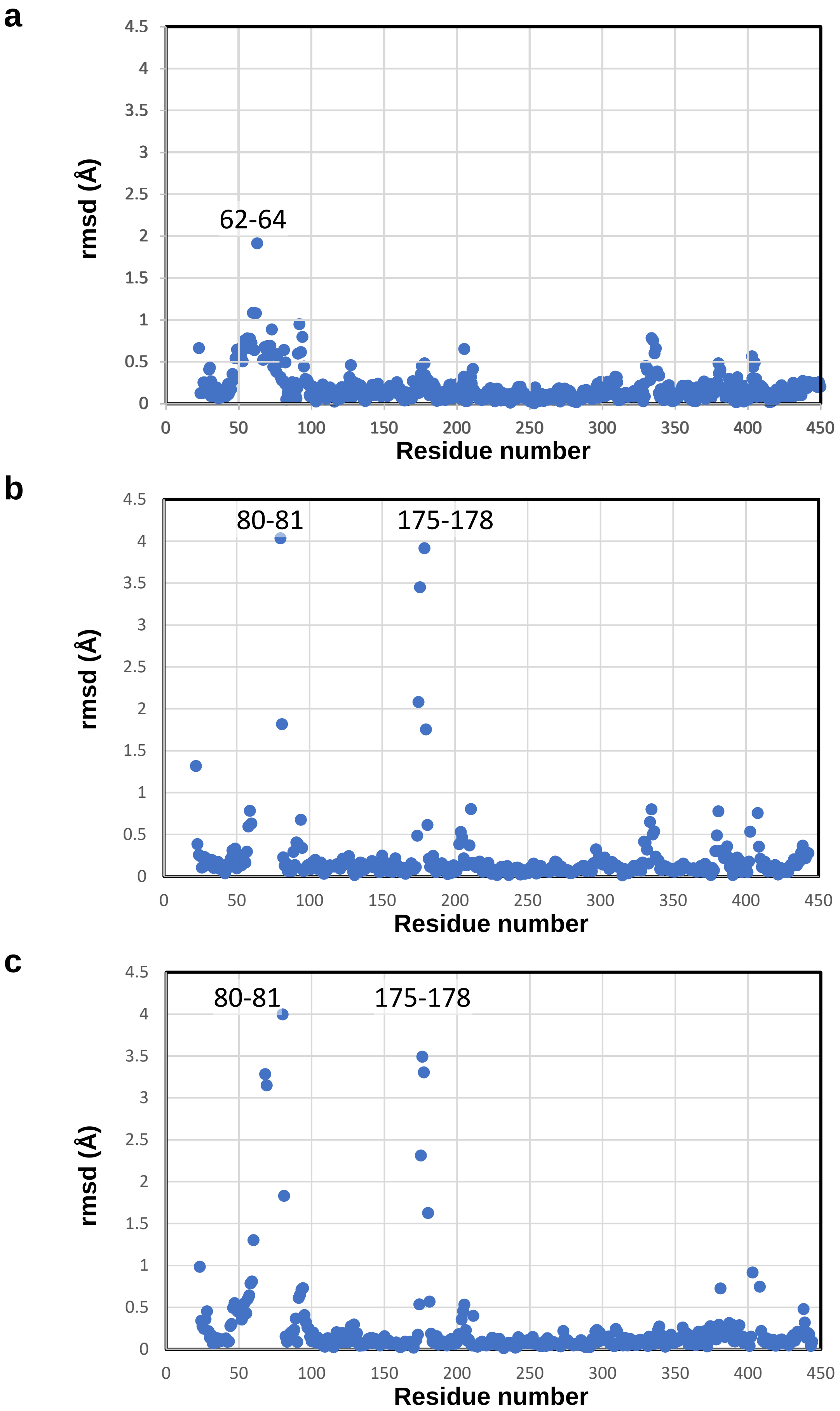


**Fig. S11 Backbone Cα atoms rmsd of RbcL. a-c**, Rmsd of RbcL between Col-0 and M309I (a), Col-0 and D397N (b), and M309I and D397N (c).

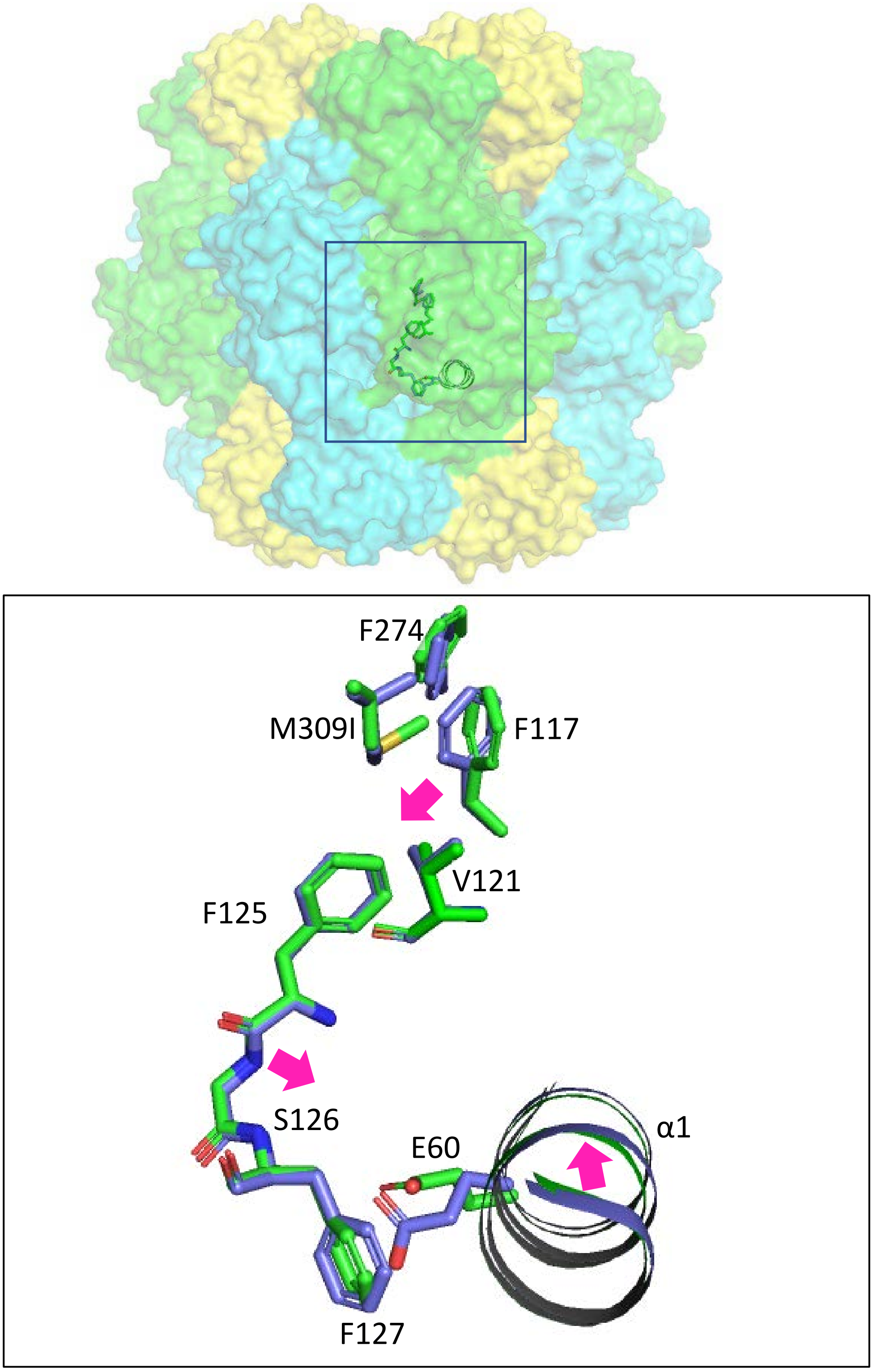


**Fig. S12 Possible mechanism of structural changes induced by the M309I substitution.** Superimposed structures of Col-0 (green) and M309I (blue) Rubisco are depicted. The overall structure of Rubisco is shown at the top, while the magnified view of the region surrounding the α1 helix is presented at the bottom. The magenta arrows indicate potential structural shifts caused by the substitution of M309 to I.

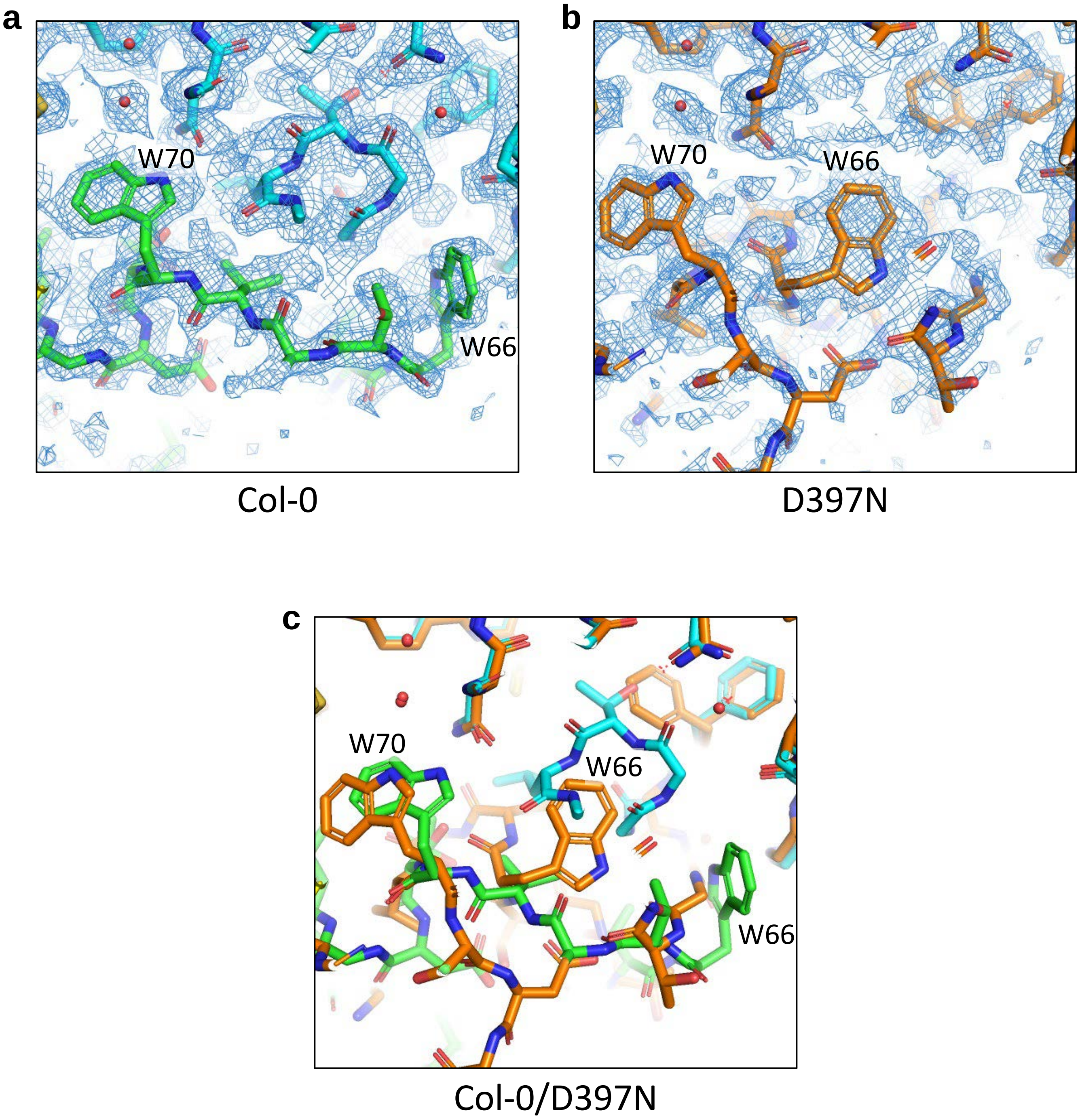


**Fig. S13 Differences in cryo-EM density maps around the 60s loop between Col-0 and D397N Rubisco. a, b**, The cryo-EM density maps contoured at 1.5 σ for Col-0 (a, green) and D397 (b, orange). **c,** Superimposition of the Col-0 and D397N Rubisco structures around the 60s loop.
